# Predictive coding yields a brain-like code for shape from motion

**DOI:** 10.64898/2026.05.13.724755

**Authors:** Yoon H. Bai, Thomas P. O’Connell, Yoni Friedman, Ani Ayvazian-Hancock, Hannah Maver, Joshua B. Tenenbaum, James DiCarlo

## Abstract

Interacting with the world relies on knowing its three-dimensional structure, yet vision receives a stream of two-dimensional images. Recovering 3D geometry from dynamic input is therefore foundational to vision, but lacks a computational account. Here we identify one, using artificial neural networks to predict human behavior and neuronal responses from the two major visual pathways in macaques. Using videos of camouflaged objects whose shape is revealed through motion, we found both pathways carry information for motion-defined surfaces, while neurons in parietal cortex most closely track what humans perceive. Only predictive coding video models—trained to fill in missing content in natural videos—reproduced both neuronal responses and human behavior. Predictive coding thus emerges as a principle that yields a brain-like geometric world model of the physical world.

## 1 Introduction

A foundational problem of vision is that the physical world has three dimensions but the eye delivers two dimensional images. Every image projected onto the retina collapses depth, and the image changes anew at every moment of motion – of objects or of the observer. Yet from this flat, variable input, we experience a relatively stable world [1] and rich enough in geometric detail to guide hand grasps and navigate complex environments. How the brain builds an internal model of the world from dynamic input remains a longstanding question that lacks a computational account.

The past decade has shown what such an answer could look like, but for a different problem. Artificial neural networks (ANNs) optimized for object recognition have become the leading image-computable models of the ventral visual pathway, one of the brain’s two major visual pathways, predicting key features of the ventral hierarchy at the level of neural spiking activity [2–4] and perceptual judgments of object category [5]. However, recognizing and labeling static images is only part of biological vision. A complete account must also explain the dorsal visual pathway, which specializes in motion cues and is also implicated in processing geometric structure for guiding action [6–9]. Here, modeling has not kept pace. Several efforts have modeled dorsal computations [10–12], yet none has reached the benchmark status that leading networks hold for the ventral stream, and the dorsal pathway lacks the neural benchmarks needed for standardized model evaluation. Closing this modeling gap calls for a dorsal analog of a behavioral goal, just as object categorization in static images facilitated investigations of ventral stream computations.

We propose recovering three-dimensional shape from motion as that goal. The recovery of three-dimensional structure from motion has been studied in perception since the kinetic depth effect [13] and in computer vision as structure-from-motion (SfM) [14, 15], providing theoretical hypotheses for biological motion-based reconstruction. At the level of local motion processing, much is known–neurons in area MT detect motion coherence and carry signatures of depth [9, 16, 17]. But recovering a coherent 3D shape demands more than local motion analysis–a requirement that has led to hypotheses of grouping local cues into a unified structure across downstream visual areas including MST and parietal regions in the dorsal pathway [18, 19]. Nonetheless, how these downstream stages transform local motion into integrated three-dimensional structure remains far less understood than the corresponding stages of the ventral hierarchy. Surface geometry from moving objects also cuts across the classical division of labor between the two streams [6, 7, 20, 21]. It is both a property of object form and a product of object motion, motivating a direct test of how each pathway contributes.

Here we address this topic by combining human psychophysics, chronic macaque electrophysiology, and a large, stimulus-computable model zoo. To isolate surface-from-motion processing, we designed rotating abstract objects rendered with identical foreground and background textures, camouflaging static boundaries and forcing the visual system to rely on motion-based grouping to recover three-dimensional structure (Figure 1). We measured human behavioral judgments on a surface-matching task using these video stimuli, and recorded from downstream regions of both pathways – inferior temporal (IT) cortex in the ventral stream and the inferior parietal lobule (IPL) in the dorsal stream – while macaques viewed the same stimuli. To reveal the computations driving neural representations, we evaluated a diverse set of stimulus-computable models spanning classical motion energy filters, supervised and self-supervised convolutional networks (CNNs) and vision transformers (ViTs), image foundation models, supervised video classification models, and predictive coding video models (i.e., models trained to predict masked content from natural video; the term as used here is defined in Section 2.4).

**Fig. 1.**
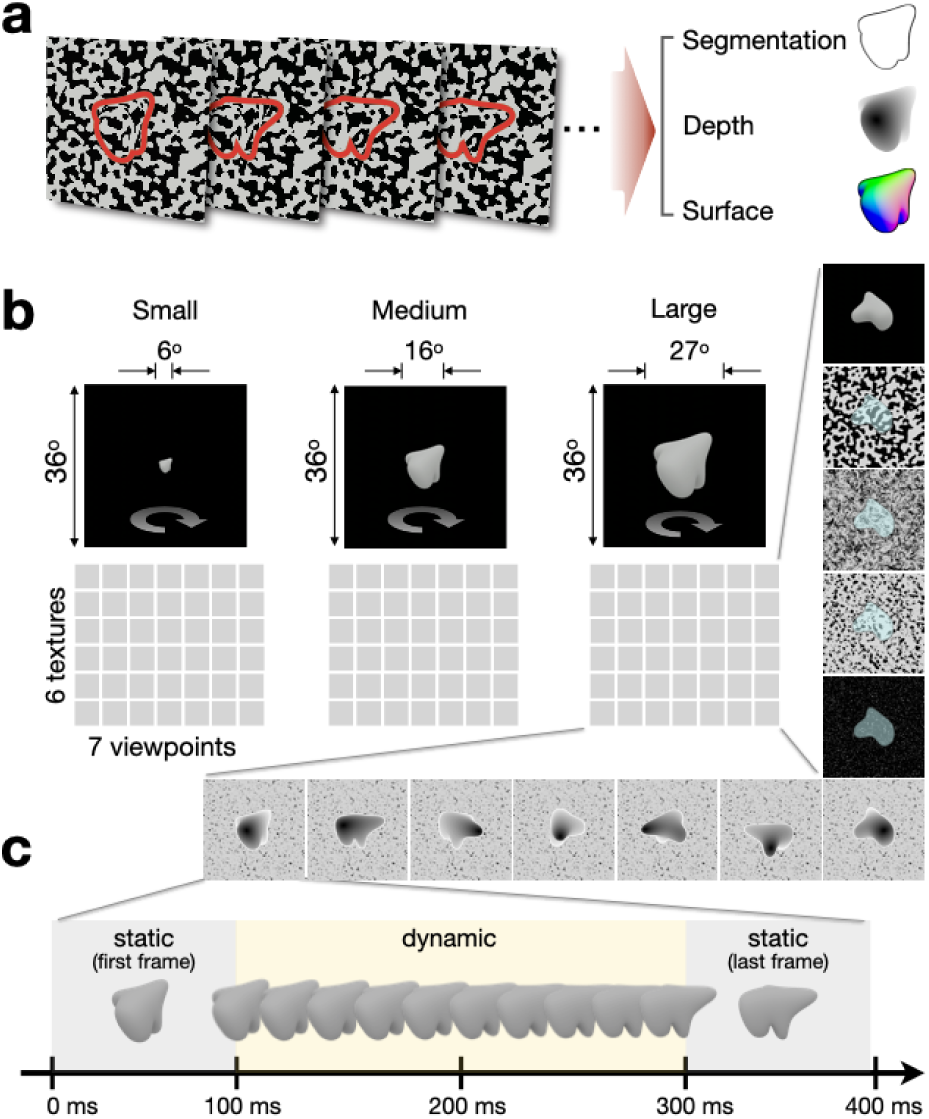
Stimulus configuration and experimental conditions. (a) **Shape-from-motion:** Surface geometry becomes perceptible when texture-masked objects undergo rotation. An abstract-shaped object is shown across four example frames, camouflaged by identical foreground-background texture. The object is outlined in red for visualization only since boundaries are difficult to discern in the absence of motion. Surface geometry can be characterized through continuous fields including segmentation masks, depth maps, and surface normals. (b) **Stimulus conditions:** Twenty unique abstract objects were presented for 400ms in video clips, across systematic variations in size (6, 16, and 27° visual angle), surface texture (five texture types plus an untextured control), and viewpoint (seven non-overlapping orientations spanning approximately 50° of rotation). Objects were centered within a 36x36° field of view. This factorial design yielded 2,520 unique video stimuli (20 objects x 3 sizes x 6 textures x 7 viewpoints). (b) **Temporal structure**: Each 400 ms video began and ended with 100 ms static periods (grey epoch) to isolate motion-driven responses from luminance transients. The intervening 200 ms dynamic period (yellow epoch) contains the rotating object; subsequent analyses focused on neural responses during this epoch.

## 2 Results

### 2.1 Humans discriminate motion-defined surfaces

3D objects that must be inferred from motion cues were designed by applying identical texture to 3D object mesh models and the backgrounds on which those object models were rendered. This effectively camouflaged the boundaries that could otherwise support object-background segmentation (Figure 1a). Under this texture-masking paradigm, the object’s surface geometry facing the viewer is largely imperceptible, but it becomes strongly perceptible as the object is rotated (see example stimuli here). Both of these psychophysical statements are quantified below.

In each video clip, the object rotated for 200ms. The moving segment was flanked by a 100ms static frame at the start and end for a total clip duration of 400ms (Figure 1b). We padded the moving segment with static frames to separate stimulus-onset and offset(luminance transients) away from object motion (rotation). This temporal structure let us examine both the object-discrimination performance and the time course over which motion information is integrated into surface geometry representations. We used identical clips for both human psychophysics and macaque neural recordings.

To quantify sensitivities to surface geometry, we asked participants (N=1063, tested online) to perform a match-to-sample task (Figure 2a, see Methods). We first presented a 400 msec video clip generated by a rotating target object followed by two rotating choice objects. One matches the same abstract object as the target object, shown rotating through the same viewpoint. The other is a lure, which is a different abstract object rotated by the same amount. Critically, the surface textures always differed between the target and the choice objects, requiring observers to rely on the three-dimensional shape or surface revealed by motion. We used this two-alternative forced choice (2AFC) paradigm as it reduces the continuous process of observing 3D surfaces into a single accuracy measurement and provides a common metric for comparing perceptual, neural, and model sensitivities in the sections that follow. As a control to isolate the contribution of motion, a separate group of participants (N=295) viewed the target as a single static frame (first frame of the same video clips), while the two choice objects continued to rotate as before. This group performed substantially worse than observers who saw the rotating target object. This confirms that human observers are highly sensitive to shape defined by differential motion patterns in our stimuli and provides a behavioral signature (patterns of matching successes and failures) to compare with neural and model task overall performance and behavioral signatures in subsequent analyses.

**Fig. 2.**
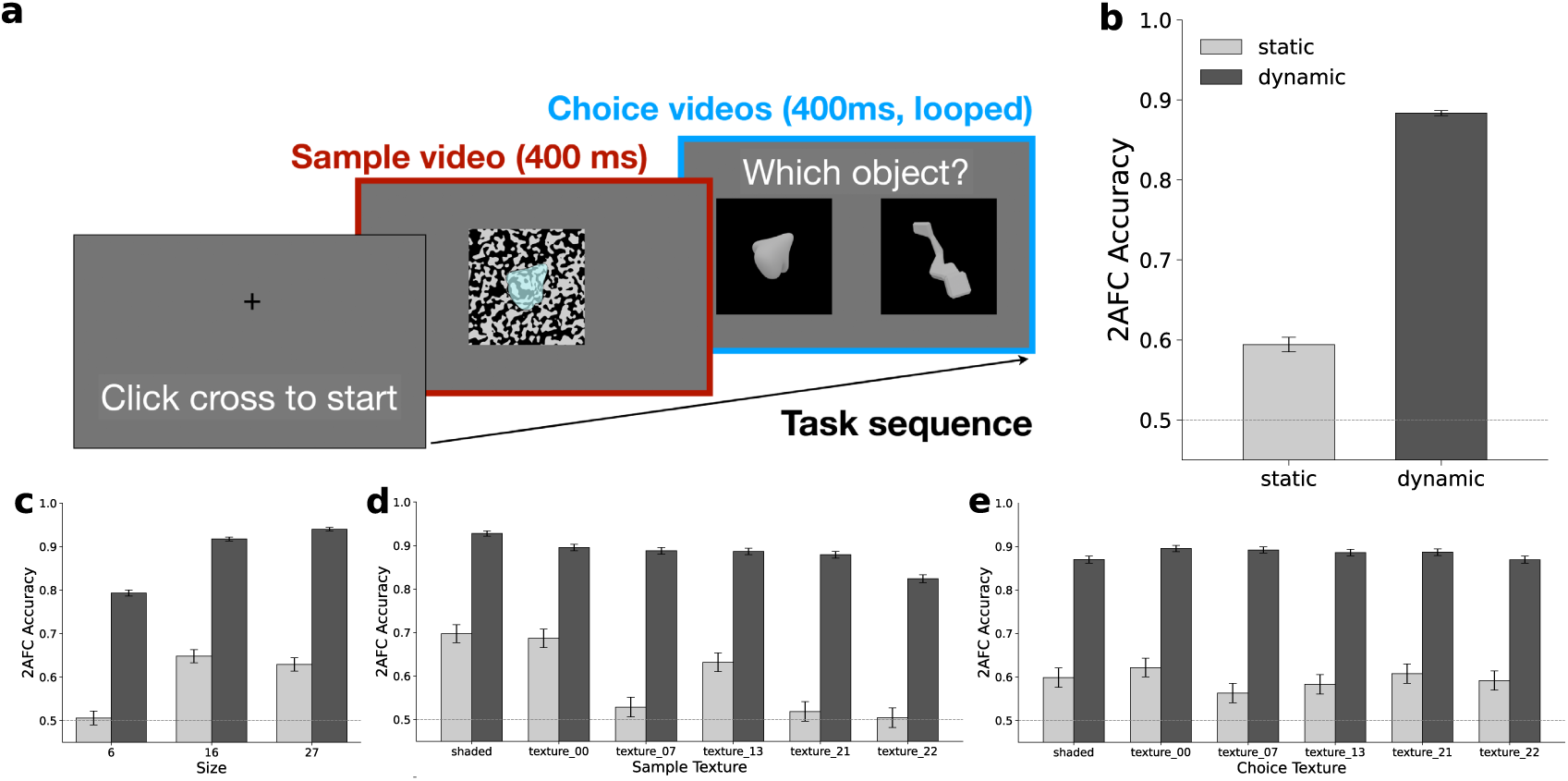
Behavioral matching of motion-defined surfaces. (a) **Task sequence:** A match-to-sample task quantifies human performance in matching motion-defined surface geometry through binary choices. Each trial begins when the observer clicks the cross sign. The sample (target) object is presented in a short video clip lasting 400 ms (red box), showing a rotating object camouflaged by a foreground-background texture (target highlighted for visualization only). The sample stimulus is followed by two choice clips (also 400ms each, looped until observer responds), and the observer selects the clip matching the previously shown object. In the static condition, the sample video was replaced with the first frame of the corresponding video, presented for the same 400ms duration, while the choice clips remained dynamic videos. (b) Humans showed substantially higher accuracy in matching motion-defined surfaces than static frames from the same videos. (c-e) This pattern holds across all experimental conditions. Error bars indicate SEM. Horizontal dotted line represents chance level performance.

### 2.2 Both pathways carry linearly decodable surface information

Humans readily discriminate motion-defined shapes even under challenging texture-masking conditions, but where in the brain is this computation performed? The classical division of labor predicts a dissociation in which motion-based computation is carried out in the dorsal stream and the resulting shape representations reside in the ventral stream. On this account, the identity of a motion-defined object should be decodable from ventral populations, which are thought to house these shape representations, but not from downstream dorsal cortex. To test this, we recorded from downstream regions of both pathways in macaque monkeys viewing the same stimuli. In the ventral pathway, we targeted anterior (aIT), central (cIT), and posterior (pIT) subregions of the inferior temporal (IT) cortex, which represents the culmination of the ventral pathway and reflects a hierarchical gradient from visual to increasingly semantic representations of object categories [22, 23]. In the dorsal pathway, we recorded from subregions of the inferior parietal lobule (IPL) – areas 7op (parietal operculum) and Tpt (temporo-parietal; upstream of area 7op) – as a counterpart to IT cortex in the ventral stream [24–28]. We used chronic arrays, as they allowed us to track the same populations across sessions and measure neural responses for the full stimulus set (2,520 videos). Achieving stable recording deep within IPL sulcal cortex (Supplemental Figure S1) was itself a methodological advance and the details are addressed in Methods (see *Target sites for electrophysiology recordings*).

To ask what object information is carried by neural population activity, we decoded a two-alternative choice directly from the recorded responses – a neural readout – using the same match-to-sample structure as the human task. On each trial, a training-free distance decoder compared the population response elicited by each choice stimulus to the response elicited by the sample (target) stimulus, and reported the choice whose response pattern was closer. Because this decoder has no fitted parameters, it provides a conservative estimate of the object information available in the population. We then applied this readout in a sliding 100 ms window (20 ms steps) to track how readout performance evolved over time (see Methods, *Applying linear decoders to neural responses across time*). An alternative approach, using logistic regression classification, yielded similar results (Supplemental figure S3). We found that decoding accuracy emerged rapidly after motion onset and was sustained throughout the dynamic period, indicating that downstream regions in both pathways (ventral: aIT; dorsal: 7op, Tpt) carry information about motion-defined surface geometry (Fig. 3b).

**Fig. 3.**
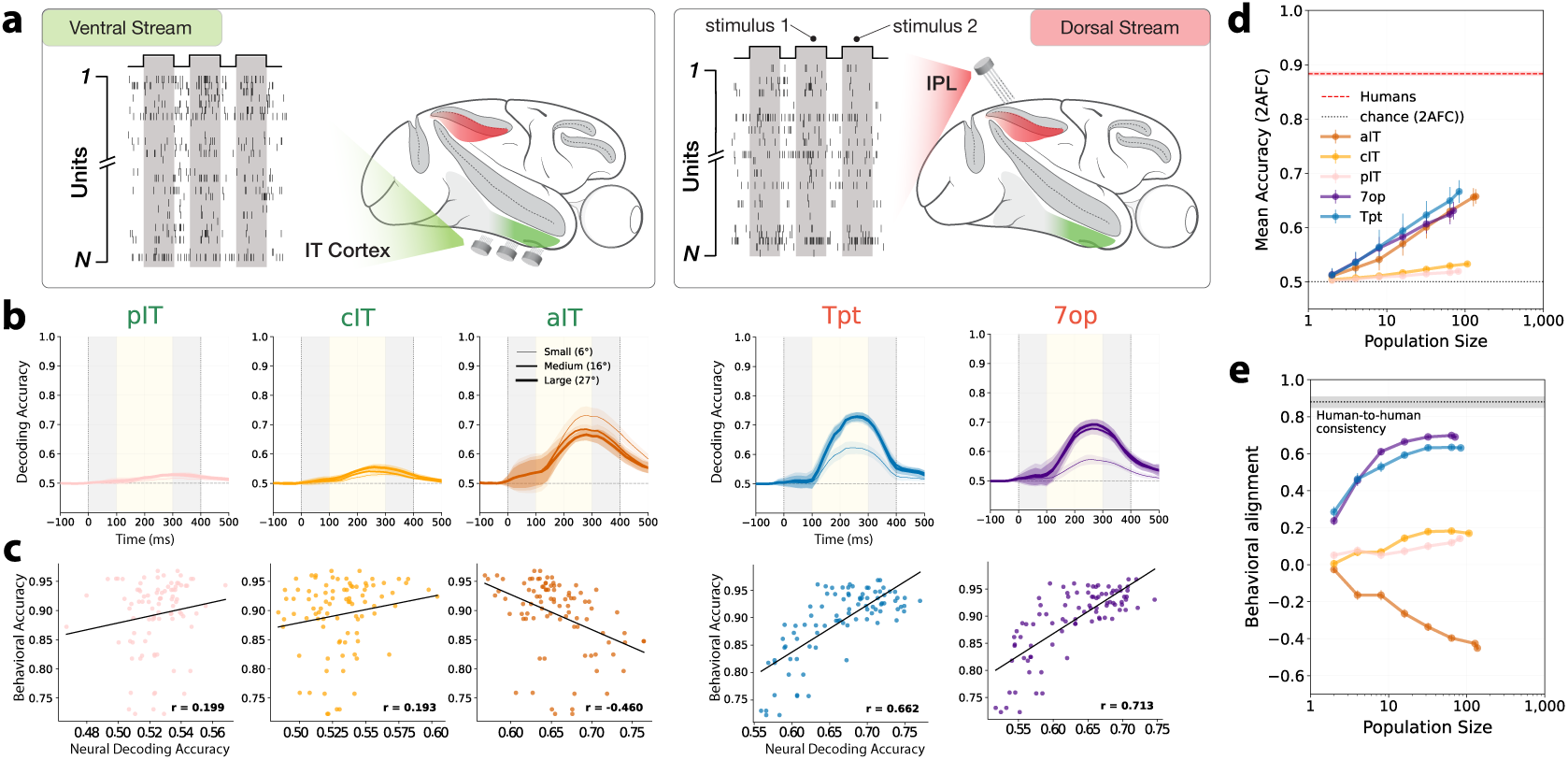
(a) **Neural recordings**. Spiking activity were recorded from both ventral and dorsal streams using chronic multi-electrode arrays. Ventral recordings targeted IT with surface arrays. Dorsal recordings targeted IPL regions buried within sulci using volumetric arrays. (b) **Neural decoding**. Downstream regions of both ventral and dorsal streams carry explicit (linearly decodable) representations of motion-defined surfaces. A single linear decoder was trained on spike rates estimated from the dynamic period (100–300ms), pooled across all three object sizes, and evaluated on held-out trials. Decoding performances are shown separately for each object size (6°, 16°, and 27°) with increasing line thickness corresponding to increasing object size. Rectangular regions in grey and yellow indicate periods of static and dynamic frames, respectively. Shaded regions indicate SEM across cross-validation folds. (c) **Neural readouts versus human accuracies**. We correlated condition-wise accuracies across the same 90 conditions tested on human participants and from neural decoding. (d) **Neural scaling**. Neural decoding performances over different numbers of resampled units. Error bars indicate SEM from resampled units; grey dotted line indicates chance performance at 0.5. (e) **Behavioral alignment versus neural scaling**. Alignment between neural readouts and behavioral performance is plotted against the number of resampled neurons. Alignment is quantified with the Spearman correlation coefficient calculated across accuracies from 90 conditions. Error bars indicate SEM from resampled units. Grey shaded band indicates 95% percentile interval of human-to-human consistency, estimated as the condition-wise split-half reliability of human condition means.

### 2.3 Linear decodes of dorsal regions track human behavioral patterns more closely than ventral readouts

Carrying linearly decodable information does not necessarily imply equivalent contributions to shape perception in dynamic scenes. For example, across the 90 behaviorally tested conditions, the condition-wise pattern of IPL decoding accuracy correlated more strongly with the corresponding pattern of human discrimination performance than did the IT readout pattern (Figure 3c). To ask whether these differences reflected population size, we examined how each quantity – overall decoding accuracy and behavioral pattern correspondence – scaled with the number of units. First, following the population-to-behavior scaling approach of [23], overall decoding accuracy grew with the number of units in both pathways but along different trajectories. IPL populations reached human-level accuracy with far fewer units than IT, indicating that dorsal representations are more efficiently organized for surface discrimination (Figure 3d). Second, scaling behavioral pattern correspondence in the same way revealed an even sharper divergence between pathways (Figure 3e). Area 7op, downstream of Tpt within IPL, showed the highest correspondence to human behavior, whereas the lowest was in aIT, the most downstream ventral region. The divergence in behavioral correspondence raises the possibility that downstream ventral regions do not represent the primary information driving perceptual judgments about surface geometry. Together, both pathways support linear readouts of motion-defined surfaces, but dorsal parietal representations are more predictive of patterns of human perceptual reports, suggesting a stronger mechanistic linkage to those reports.

### 2.4 Computational models that aim to estimate 3D object shape from dynamic input

What computations transform motion cues into neural representations of surface geometry? To address this question, we draw from the past decade of success in demonstrating strong alignment between the internal representations of some artificial neural network (ANN) models and ventral visual stream representations. Previous benchmarking has established some CNN models as the leading stimulus-computable models of the mechanisms operation along the ventral visual pathway [4], but stimulus-computable models of the dorsal pathway remain unclear, in part because systematic comparison of behavioral and neural responses to video stimuli has lagged behind responses to images. To ask whether video processing networks have internal functional alignment with IPL neural representations, we evaluated a large model zoo including classical models of motion processing with prior neural validation [10, 18, 29] (Figure 4; Motion-energy) and a diverse set of ANNs spanning image or video pretraining.

**Fig. 4.**
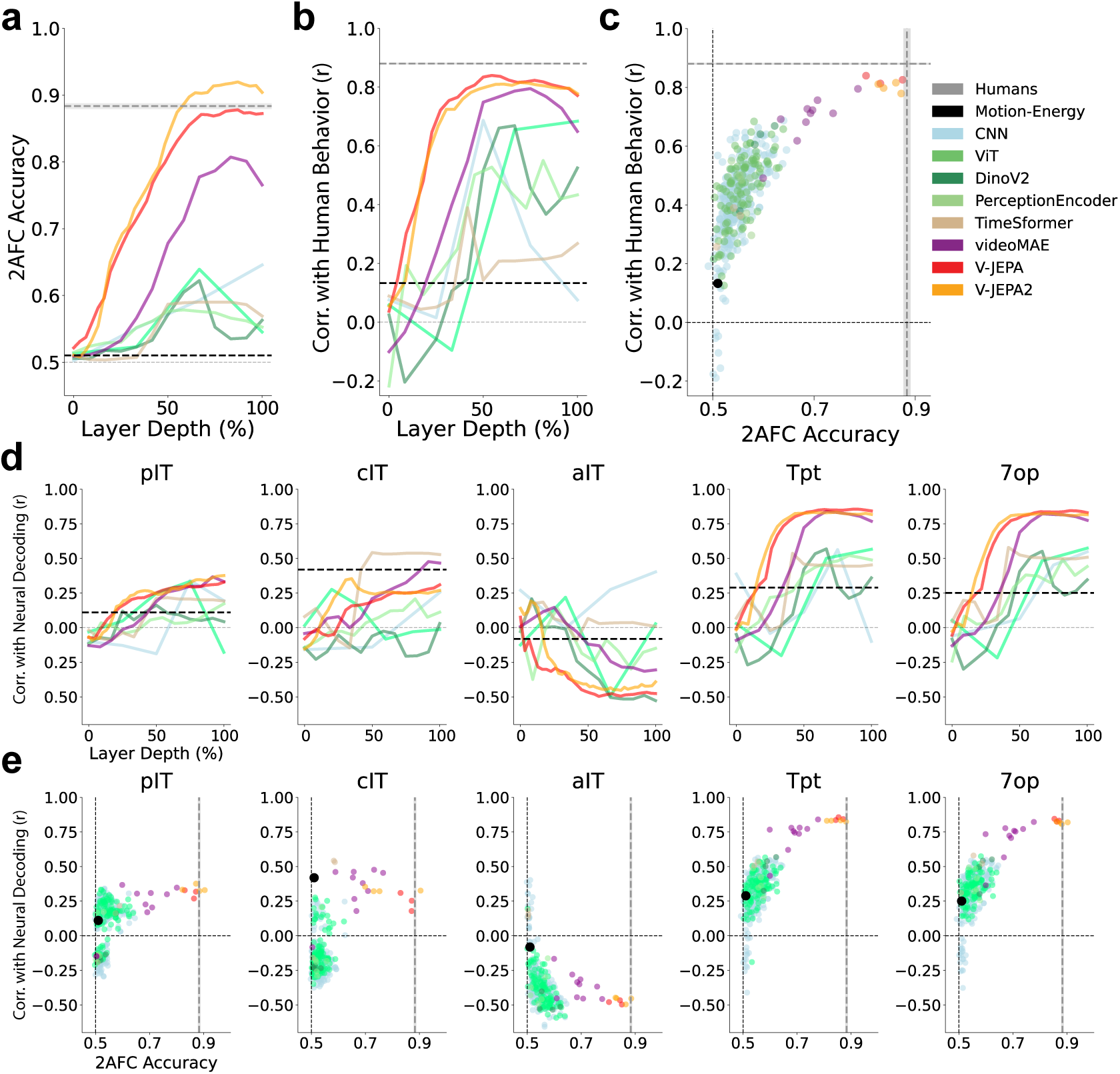
(a) **Model accuracy** on the surface-matching task as a function of layer depth. For each model class, the highest accuracy across models within that class is shown at each layer. (b) **Model consistency** with condition-wise human 2AFC performance as a function of layer depth, using the same layer-wise selection procedure as in (a). (c) **Model accuracy versus behavioral consistency** for each model’s best-performing layer (selected by peak behavioral consistency). Predictive coding video models occupy the highest accuracy–consistency regime. (d) **Model consistency to condition-wise neural decoding performance** on 2AFC task across layers. Predictive coding video models showed the highest consistency, especially to dorsal regions. (e) **Model performance x neural decoding consistency for best layer per model.** Each model’s best layer is defined by highest absolute value neural decoding consistency).

Our model set spans two axes that separate the contributions of input modality and training objective (Methods, Model zoo). Along one axis, models progress from static-image to video input; along the other, from supervised classification to self-supervised prediction. Image models include supervised CNNs and ViTs [30], self-supervised DINOv2 [31], and PerceptionEncoder, a contrastive vision–language foundation model [32], none of which learn explicitly from temporal structure. Video models include supervised TimeSformer [33] and the self-supervised VideoMAE [34] and V-JEPA [35, 36]. We refer to V-JEPA and VideoMAE collectively as *predictive coding video models*, to emphasize a shared training strategy — learning representations by predicting withheld sensory content, including future time steps from past context. Our terminology is broader than the classical predictive coding framework in neuroscience [37, 38], in which top-down predictions are compared against bottom-up input and prediction errors propagate downstream. In summary, the progression in our model zoo lets us ask which computational paradigm best approximates the neural mechanisms that extract surface geometry from motion.

### 2.5 Predictive coding video models exhibit best surface-matching performance, with alignment to human behavior and IPL readouts

Running each model through the same match-to-sample decoding pipeline used for neural decoding (Methods), we found that predictive coding video models reached human-level accuracy. For predictive coding video models, both overall accuracy (Figure 4a) and condition-wise correspondence with human behavioral patterns (Figure 4b) were highest at middle-to-late layers, consistent with surface-from-motion features building up at later processing stages. To directly compare the model families, we summarized the accuracy and behavioral consistency of each model using its best-performing layer (Figure 4c). Predictive coding video models (V-JEPA and VideoMAEs) consistently achieved the highest accuracies and closest alignment with human behavioral patterns.

Finally, we measured the consistency between model readouts and neural readouts from different cortical regions. We observe that the middle-to-late layers of predictive coding video models show high consistency with neural readouts from IPL (Figure 4d,e). While predictive coding video models showed the highest consistency with neural readouts from upstream ventral regions (pIT, cIT), this pattern reversed in the most downstream region (aIT), where CNN architectures achieved the highest model-neural consistency and video models showed consistent negative correlations. The negative correlations between predictive coding video models’ readouts and aIT readouts is consistent with the negative correlation between human behavior and aIT readouts. This dissociation suggests that the computations underlying motion-based surface estimation may be differentially distributed across visual pathways, with high level dorsal regions more involved in the estimation of surface geometry from dynamic inputs and high level ventral regions more involved in estimating semantic information from static images [21–23]. To summarize, predictive coding video models, particularly V-JEPA models, achieve human-level accuracy on 3D object shape estimation from dynamic input, approach the human noise ceiling for consistency with condition-wise human performance, and their internal features show high condition-wise correspondence with IPL (dorsal stream) neural population readouts – but not IT (ventral stream) readouts.

### 2.6 Encoding analyses: computations transforming pixels to representations of surface-from-motion

Across our model zoo, distinguishing objects defined by surface geometry showed that predictive coding video models best carried geometric information persisting through camouflaging texture. Furthermore, their alignment with human performance patterns hints at shared visual processing algorithms. Such behavioral alignment, however, does not by itself establish whether models and neurons transform visual input into internal representations in similar ways[39, 40]. We therefore asked whether models and neurons transform visual input into latent representations in a similar fashion, using two complementary encoding analyses, each applied to every layer of each model’s encoder. First, univariate prediction of individual neural unit responses from model internal feature spaces, which measures shared feature selectivity between model units and single neurons. Second, multivariate prediction of representational geometry, which captures the population structure that could be utilized for downstream readout [41, 42].

#### 2.6.1 Univariate analysis – predicting individual unit responses

We mapped model features onto individual neurons using Partial Least Squares (PLS) regression, extending an approach validated on static images to dynamic video [3, 4] (see Methods and Appendix S5.1 for details). Across ventral and dorsal regions, video models outperformed image-based models, with VJEPA-2 achieving the highest neural response prediction score (Fig. 5a). For top-performing video models, middle-to-late model layers yielded the highest neural prediction score, consistently across all brain regions. Comparing each model’s mean predictivity with its 2AFC accuracy (Fig. 5b), models with higher predictivity generally performed better, particularly in dorsal regions.

**Fig. 5.**
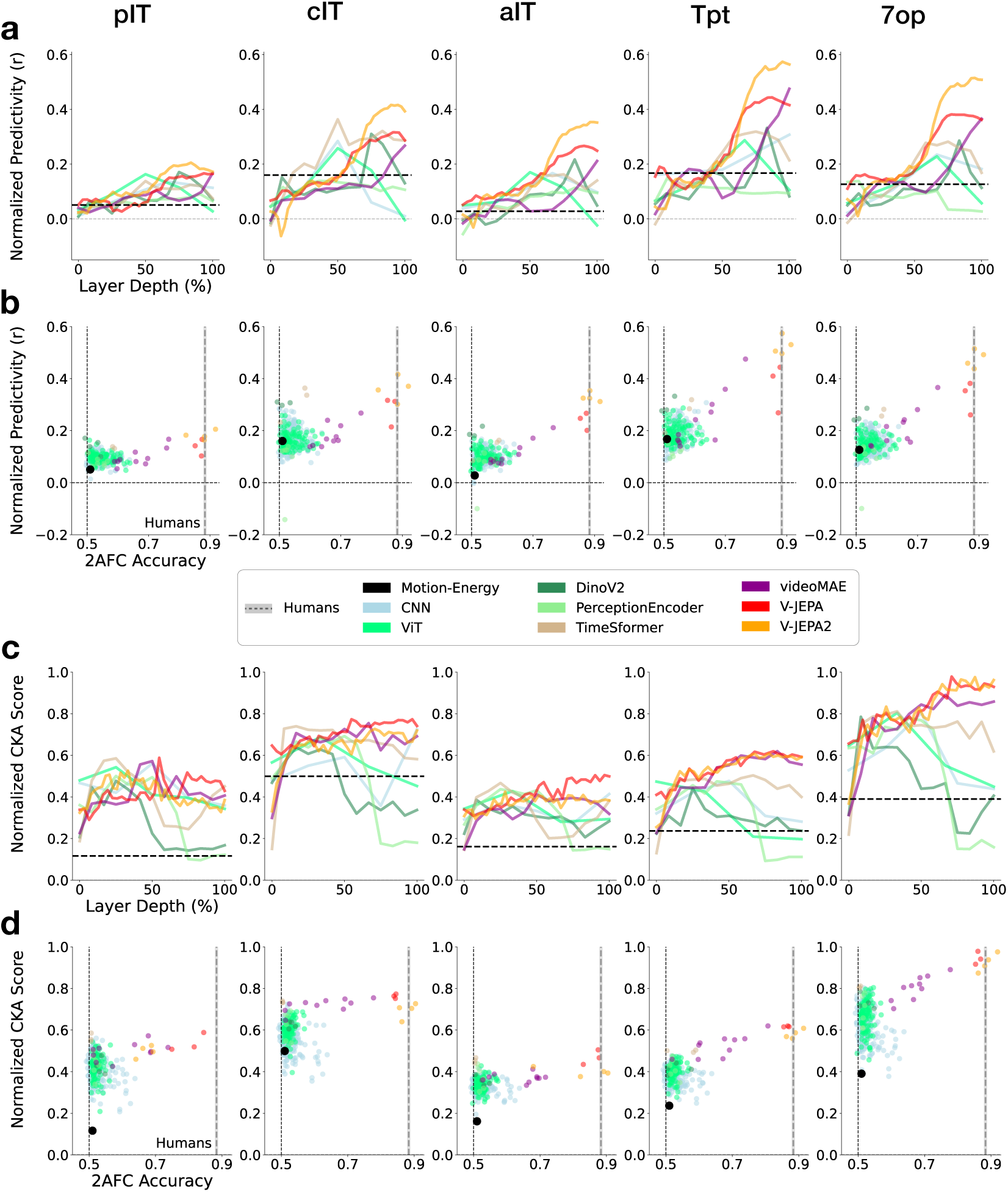
Encoding comparisons between models and neurons recorded from downstream regions of the ventral (pIT, cIT, aIT) and dorsal (Tpt, 7op) visual pathways. Each cortical region is shown in the same column across panels. **Single-unit predictions**: (a) Neural responses predicted from linear combinations of model features, at each model layers. The motion-energy model lacks layers and is shown as a black horizontal dotted line. (b) Predictivity versus 2AFC accuracy for the best (highest-predictivity) layer from each model. **Predicting population-geometry**: (c) We quantified the similarity between predicted and held-out neural population geometry with Centered Kernel Alignment (CKA), at each model layer. Layer-wise alignment is shown for the best-performing model in each model class, normalized to the noise ceiling. The motion-energy model lacks layers and is shown as a black horizontal dotted line. (d) Normalized population alignment versus 2AFC accuracy for the best (highest-alignment) layer from each model. The motion-energy model is marked with a black dot.

#### 2.6.2 Multivariate analysis – predicting population responses

Models with similar single-unit predictivity, or explained variance, can nonetheless arrange stimuli very differently in their representation spaces, and the pairwise distances that matter for decoding may be invisible in unit-by-unit predictions (see Supplemental Figure S4). We therefore quantified population-level alignment with Centered Kernel Alignment (CKA, Supplement S5.2), which emphasizes the high-variance directions dominating a population’s response and tracks how well one population can be linearly decoded from the other – making it well suited to assessing geometric structure exploitable by downstream linear readouts [41]. We applied the same hold-out policy and analyzed dynamic-period responses as above.

Predictions from multivariate analyses both corroborated and extended the univariate findings (Fig. 5 c,d). First, predictive coding video models showed the highest geometric alignment with neural populations in both IT cortex and IPL. Second, their advantage over PerceptionEncoder – a vision-language model (VLM) trained at far larger scale^1^ than any other model evaluated here–was more pronounced for population geometry than for single-unit predictivity. Third, the two analyses revealed distinct layer-wise profiles – in predictive coding video models, population alignment rose earlier in the hierarchy than the late-stage rise seen with single units. A time-resolved analysis yielded the same rank ordering, confirming robustness to temporal aggregation (Supplementary Fig. S6).

Overall, both univariate and multivariate analyses converged on predictive coding video models as top performers and showed that self-supervised video models trained on relatively modest datasets outperform alternative models trained at larger scale. Furthermore, the two approaches yielded complementary insights. Univariate analysis showed that video models–including TimeSformer–predict individual responses well, reflecting shared feature selectivity, whereas multivariate analysis showed that only V-JEPA and VideoMAE preserve the population geometry with high fidelity, which could be usable at subsequent stages in the brain.

### 2.7 Establishing leading models of high level areas of the dorsal pathway

The preceding sections evaluated models of surface-from-motion processing along four complementary measures: behavioral accuracy, behavioral consistency with human observers, predictivity of individual neural sites, and neural population-level representational geometry. Across these measures, the convergence of behavioral patterns with neural readouts in IPL identifies this dorsal region as more closely linked to human judgments of surface geometry than the IT ventral visual region.

Identifying a candidate model of these computations require more than fitting recorded responses. A stronger test asks whether a model also produces read-outs that match those of the recorded population, on stimuli it was never fit to. We therefore combined our encoding and decoding analyses into a joint test on held-out stimuli – a criterion that constrains candidate mechanisms more tightly than either analysis alone. For each region we built synthetic surrogate populations by mapping model features onto recorded responses (PLS and CKA; Methods), then read-out match-to-sample choices from those surrogates on held-out objects and textures. Each surrogate’s generalization was scored across two axes, as shown in Figure 6: encoding fidelity (y-axis) — how well it predicts held-out neural responses, and read-out similarity (x-axis) — how closely its condition-wise decoding matches the recorded population on unseen objects and surface textures.

**Fig. 6.**
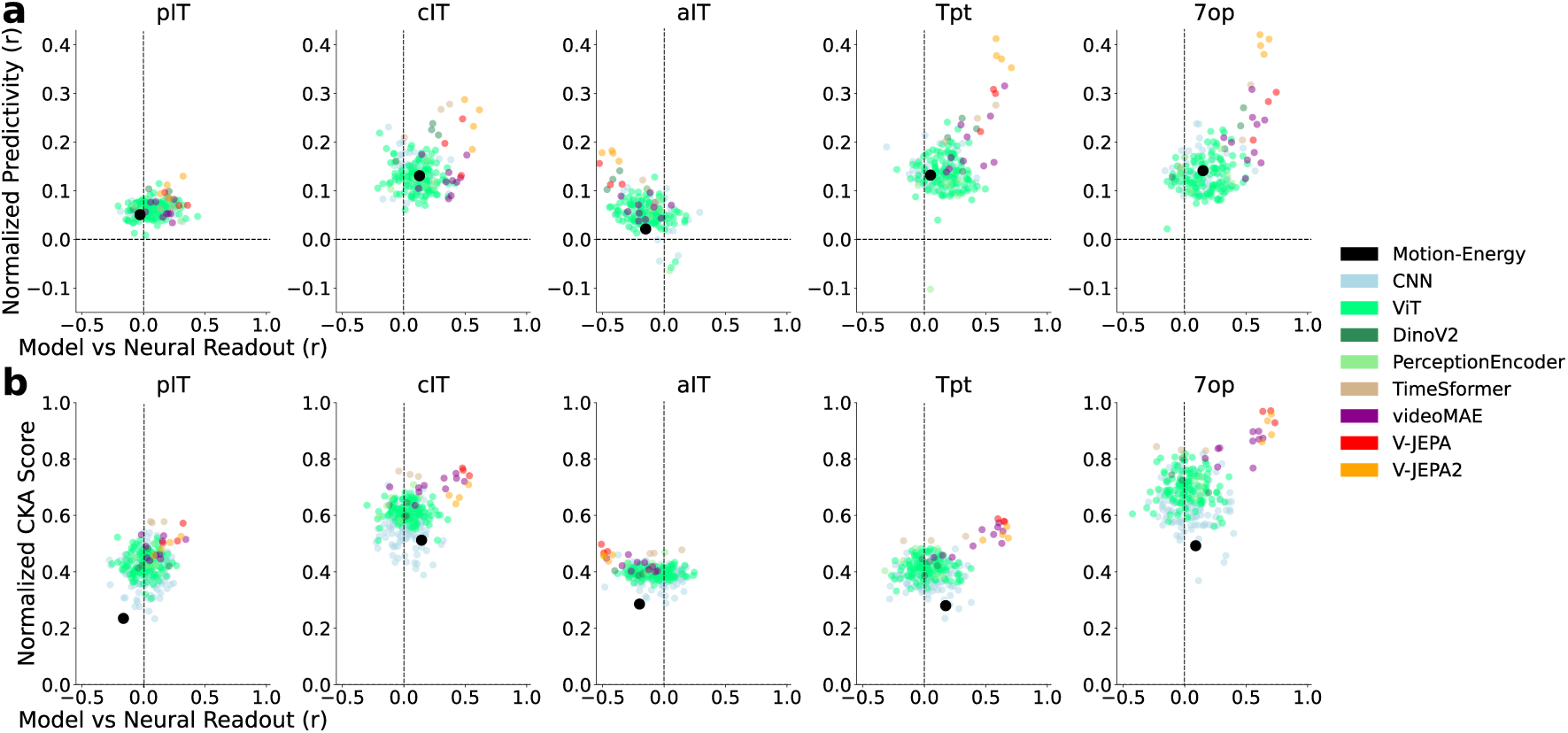
Generalizing neural encoding and decoding from synthetic populations. (a) Synthetic populations generated unit-by-unit from stimulus-computable models via PLS regression. Each column corresponds to a cortical region (as in Figure 5); points are color-coded by model class. The x-axis shows readout similarity between model and biological neurons, defined as the Spearman correlation between readout accuracies across held-out conditions. The y-axis shows the encoding correlation between PLS-mapped synthetic responses and recorded neural responses, averaged across held-out conditions. (b) Synthetic populations generated at the population level by matching coordinated activity across neurons (CKA). Plotting conventions as in (a). Raw plots without normalization are shown in Supplement S7.

On encoding fidelity alone, predictive coding video models best matched neural responses in both pathways. However, jointly evaluating the two axes revealed a pathway dissociation. In dorsal regions these models generalized on both axes, whereas in ventral regions the same models achieved high encoding fidelity yet produced readouts that diverged sharply from the recorded population. Two accounts could explain this dissociation. First, the ventral pathway may implement computational strategies for surface-from-motion that our model zoo does not capture. Alternatively, ventral regions may carry surface-geometry information – consistent with their above-chance decoding (Section 2.2) – that is not organized for the linear match-to-sample readout our surrogates reproduce. Distinguishing these accounts will require targeted perturbation, either through invasive or non-invasive methods, and we leave this to future work.

Together, our results support a specific computational insight. Among the models we tested, predictive coding objectives (predicting masked stimulus features or embedded features across space and time) produced the closest alignment with downstream dorsal responses and their read-out patterns. This advantage was specific to dorsal regions—the same leading models achieved comparable encoding fidelity with ventral neurons but diverged markedly in their condition-wise read-out similarity. We therefore take these models as new baseline hypotheses for how the visual system estimates surface geometry from dynamic input. One interpretation of why these objectives succeed is that predicting masked or future structure favors latent representations that preserve statistical regularities associated to geometric structure while discounting nuisance variables such as surface texture — a conjecture our results motivate but do not test directly.

## 3 Discussion

We investigated how the primate brain recovers three-dimensional shape from motion by combining human psychophysics, macaque neurophysiology, and computational modeling. Using texture-masked stimuli that isolate motion-based surface perception – challenging for computer vision models yet readily perceived by humans – we report three principal findings.

First, both visual pathways contain explicit (linearly decodable) surface geometry representations. Contrary to strict functional segregation, downstream regions in both dorsal (IPL) and ventral (IT cortex) pathways support robust decoding of alternative 3D shapes for motion-defined shapes. This finding challenges the classical view that motion processing engages primarily dorsal structures while shape representations reside predominantly in the ventral visual pathway. Instead, our results indicate that surface geometry information is distributed across both streams.

Second, high level (IPL) dorsal neural representations show preferential correspondence to surface geometry and behavior. Although both pathways carry shape information, IPL population linear readouts exhibited stronger correspondence to human behavioral patterns than IT populations. This asymmetry suggests that IPL representations are more directly linked to perceptual decisions about surface geometry, even though shape information is present in both pathways. Whether this reflects a causal contribution of IPL to these judgments, or a shared upstream source that IPL tracks more faithfully, cannot be distinguished by our correlational measurements.

Third, our investigation identifies a candidate baseline model for dynamic visual processing in the dorsal pathway. Motion energy models capture local motion processing in V1 and MT [10, 29], but do not extend to parietal representation of surface geometry. More recent models optimized for action recognition or self-motion estimation [11, 12] reach beyond local motion, but have not accounted for downstream parietal representations of three-dimensional structure. Among the diverse models evaluated here, predictive coding video models (V-JEPA and VideoMAE) achieved the highest alignment with neural activity in both pathways, with preferential correspondence to dorsal regions, and the closest condition-wise correspondence to human behavioral patterns. This preferential alignment with downstream dorsal areas positions predictive coding video models as the current leading stimulus-computable models of cortical processing in at least high level stages of the dorsal visual hierarchy.

### Toward the brain’s physical world model

Beyond establishing a baseline, our findings suggest a new perspective of what the dorsal pathway is for, and how to model it. On the classical view, the dorsal stream is a motion-processing system. Our results are more consistent with a stronger proposal, in which downstream dorsal processes underpins a predictive geometry engine, maintaining representations of the three-dimensional structure of the environment that are learned, and perhaps continuously updated, through prediction of incoming sensory input. This view connects the physiology reported here to predictive coding frameworks in neuroscience, which propose that the brain learns internal models to anticipate its sensory stream [1, 37], and speaks to a central ambition of machine learning–building world models that capture how the physical environment is structured and how it evolves.

Our results suggests that predictive objectives on natural video are a productive path toward representations with human-like geometric competence—and that neural and behavioral benchmarks like the one introduced here can adjudicate among candidate models in a way that task performance alone cannot. For neuroscience, our work delivers the dorsal pathway’s first downstream stimulus-computable baseline, a falsifiable starting hypothesis for the mechanisms by which cortex converts dynamic images into actionable geometry. One interpretation of why predictive objectives succeed is that predicting masked or future structure favors latent representations that preserve the regularities of geometric structure while discounting nuisance variables such as surface texture—a conjecture our results motivate but do not test directly.

We emphasize that our findings provide a foundation, far from a finished structure. A physical world model, in the full sense, maintains representations of persistent objects across occlusion, anticipates the consequences of forces and contacts, and conditions its predictions on the agent’s own actions. Our study establishes the first pillar—that the geometric representations underlying 3D percepts in downstream dorsal cortex are captured by predictive learning—and provides the recording foothold (chronic access to IPL across thousands of naturalistic video conditions) and the benchmark against which those richer capacities can now be tested. Extending this framework to viewpoint-invariant shape, self-motion, physical dynamics, and action-conditioned prediction is, in our view, a promising route to a mechanistic account of the brain’s model of the physical world.

## 4 Methods

### Behavioral task design

Measuring perceptual sensitivity to surface geometry poses an experimental challenge: the physical structure of an object—characterized by depth, surface normals, and curvature—varies continuously across space rather than falling into discrete categories (Figure1a). We addressed this challenge using a match-to-sample task, embedded in a 2AFC framework. On each trial, observers viewed a sample video (400 ms, 60 fps) depicting a rotating, texture-masked object, followed by two choice videos – also 400 ms each, looped until a decision was made (Figure 2a). One choice depicted the same object as the sample and the other a different object; surface textures were always swapped between sample and choice stimuli, ensuring that correct responses required extracting shape from motion cues rather than matching low-level image features. This design grounds behavioral measurement in signal detection theory [43], enabling well-characterized metrics of perceptual sensitivity.

Our approach departs from classical psychophysical paradigms such as orientation discrimination or contrast detection, in which experimenters systematically vary a physical parameter (e.g., angular difference, stimulus contrast) to map stimulus manipulations onto perceptual thresholds. No analogous metric exists for the perceptual dissimilarity between arbitrary 3D shapes, precluding precise a priori control of task difficulty. We therefore adopted a broad-sampling strategy: rather than titrating difficulty along a single dimension, we constructed a factorial design by varying multiple dimensions of the stimulus set.

We incorporated 20 unique abstract objects, presented across three sizes (6°, 16°, and 27° visual angle), six texture conditions (five surface texture plus an untextured control), and seven non-overlapping viewpoints spanning approximately 50° of rotation. This factorial design yielded 2,520 unique video stimuli, creating a multidimensional parameter space. In the match-to-sample task, however, the full combinatorial space of object pairs, sizes, textures, and viewpoints was prohibitively large. We therefore sub-sampled 90 unique conditions based on object size, sample texture, and choice texture, then randomly sampled across object pairs and viewpoints within each condition. This design allows task difficulty to emerge empirically from the stimulus space rather than being imposed by assumption.

We collected human behavioral data on the same task sequence, using two types of stimulus versions: one with video stimuli, and another with static frames (from the same videos used before). First, using video stimuli, behavioral data were collected from online participants (N=1063) who completed the task in a web browser (Figure 2a). Human observers performed well above chance across all 90 conditions (Figure 2b, dynamic). On the other hand, performance dropped substantially when the same objects were presented as static images, confirming that observers (N=295; Figure 2b, static) were more sensitive to motion-defined surface geometry rather than residual static cues from texture boundaries. Condition-wise consistency with human behavioral patterns from the 90 unique conditions served as our primary dependent variable for measuring the behavioral alignment of neural responses and model predictions throughout our analyses.

### Target sites for electrophysiology recordings

We recorded extracellular activity using chronic implants in both the ventral and dorsal visual pathways, targeting IT cortex and the IPL, respectively. These regions occupy analogous positions within their respective pathways—downstream hierarchical levels where visual neurons become increasingly multimodal, interfacing extensively with non-visual areas. IT cortex, while central to object recognition and category representation, connects extensively with regions processing semantic and mnemonic information [44–46]. Similarly, the IPL is implicated in processing higher-order motion-based visual information while also exhibiting strong responsiveness to somatosensory inputs and auditory cues, reflecting a multimodal integration profile that mirrors IT cortex’s interface with semantic networks [26–28]. This parallel organization motivated our selection of IT cortex and the IPL as counterparts in the ventral and dorsal pathways.

In the ventral pathway, we recorded from inferior temporal (IT) cortex. Three Blackrock Utah arrays (96 channels each) were implanted on the cortical surface targeting anterior (aIT), central (cIT), and posterior (pIT) inferior temporal cortex. In one male rhesus macaque (Macaca mulatta), we collected IT responses in the left hemisphere and repeated the experiment in the right hemisphere to verify our results. Across both hemispheres, all arrays were placed on the cortical surface ventral to the superior temporal sulcus (STS). Approximate stereotactic coordinates were defined with a David Kopf macaque stereotax and guided by macaque atlases [24, 25] (relative to ear bar zero, EBZ): (a) aIT: +12 mm rostral, +2 mm dorsal; (b) cIT: +7 mm rostral, +4 mm dorsal; and (c) pIT: +2 mm rostral, +6 mm dorsal.

In the dorsal pathway, a single Modular Bionics N-Form array was implanted to simultaneously target two subregions of the inferior parietal lobule (IPL): area 7op and area Tpt. Because both regions lie within sulcal cortex at different depths, we customized the shank lengths within the same array—eight shanks (64 channels) were fabricated at 11 mm to reach the fundus of 7op at the confluence of the intraparietal and lateral sulci, while an additional eight shanks (64 channels) were fabricated at 6 mm to target Tpt at a comparatively shallower depth. This configuration enabled simultaneous chronic recordings from two dorsal subregions with a single surgical insertion. We recorded from two animals; the N-Form array was implanted in the left hemisphere of the first animal and the right hemisphere of the second. Target regions within the IPL were confirmed with histological analyses in one animal (see S1). To reach regions buried in the sulcus with precision, we added two components to the implantation procedure used for implanting arrays in the ventral stream (IT cortex). First, we used CortexPlore (by cortEXplore GmbH), a surgical guidance system, to position the array along pre-planned trajectories derived from structural (MRI and CT) scans. Second, we used a NeuralGlider Inserter (by Acutated Neuroscience, Inc.) to reduce cortical tissue dimpling and aid array insertion via ultrasonic micro-vibration.

### Recordings from IT cortex (ventral visual pathway)

Recordings were obtained from both hemispheres across two rounds of recording sessions per hemisphere (4 sessions total per region), yielding populations of visually driven neurons in three subregions along the posterior-to-anterior axis: posterior IT (pIT; n = 82 units), central IT (cIT; n = 107 units), and anterior IT (aIT; n = 137 units). During each session, the animal performed passive fixation while 2,520 surface-from-motion video stimuli were presented 400ms, and spatially centered to the center of the population receptive fields ( fovea, see S2). Peri-stimulus time histograms (PSTHs) were computed in 10 ms bins spanning -200 to 600 ms relative to stimulus onset (80 time bins). To increase population size, units from all four recording sessions were aggregated by concatenation along the unit dimension, with up to 24 repeated presentations per stimulus. For sessions with fewer than 24 repeats, missing entries were padded with *NaN* values, and the mean response was estimated across available repeats (minimum of 20 repeats).

### Recording from IPL (dorsal visual pathway)

Recordings were collected from two male rhesus macaques using chronically implanted in the inferior parietal lobule (IPL), a downstream region of the dorsal visual pathway. Two regions were targeted: area 7op (parietal operculum; n = 71 units) and area Tpt (temporo-parietal area; n = 84 units). For each monkey, recordings were collected over two rounds of sessions. During each session, the animals performed passive fixation while 2,520 surface-from-motion video stimuli were presented 400ms, and spatially centered to the center of the population receptive fields (parafoveal, see S2). PSTHs were computed with the same temporal parameters as the ventral recordings (10 ms bins, -200 to 600 ms, 80 time bins). Units from two recording rounds were aggregated, with up to 26 repeated presentations per stimulus (minimum of 20 repeats). As with the ventral regions, the mean spiking response was estimated across all available repeats.

### Extracellular voltage preprocessing

Raw extracellular voltages were acquired using the Intan RHD2000 recording system at a sampling rate of 30 kHz. Because the full stimulus set (2,520 videos) required presentation across multiple recording sessions spanning days to weeks, we developed a stitching procedure to concatenate raw voltage traces from all sessions into a single continuous binary file per cortical region. Each session’s sample count was verified against the corresponding timestamp record to ensure sample-level integrity, and cumulative indexing was maintained to preserve the temporal registration of each session within the aggregate trace. This stitching step produced one *C×T* matrix per region, where *C* is the number of recording channels and *T* is the total number of samples across all recording sessions, serving as the input to spike sorting. Spike sorting was performed using Kilosort 2.0 [47, 48] on the stitched binary files. Region-specific channel maps reflecting the physical geometry of each array were provided to Kilosort. Sorted results were exported to the Phy template, a graphical user interface, for visual inspection.

We applied no additional data inclusion criteria beyond the default quality metrics provided by Kilosort. All units were retained for analysis provided that their spike waveforms exhibited biophysically plausible extracellular signatures (e.g., characteristic negative–positive deflection shape and consistent amplitude). Units were excluded only when waveform inspection revealed clear mechanical or electrical noise artifacts.

### Distance-based linear decoder

To assess the discriminability of object representations in neural population activity, we employed a distance-based match-to-sample (M2S) decoder. On each trial, the decoder was presented with a sample stimulus defined by a particular object identity, texture, viewpoint, and size. The decoder compared the neural population response to the *sample* against two alternatives: a *choice* stimulus depicting the same object in a different texture and same viewpoint, and a *lure* stimulus depicting a different object rendered in the same texture as the choice. The decoder classified the sample as the object whose population response was closest in cosine distance:

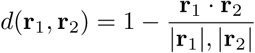

where **r**_1_ and **r**_2_ are population response vectors (firing rates across all recorded units). The decoder selected the choice over the lure when *d*(**r**_sample_, **r**_choice_) *< d*(**r**_sample_, **r**_lure_), yielding a two-alternative forced-choice accuracy for each trial.

### Logistic regression classifier

As an alternative readout, we trained an L2-regularized logistic regression classifier to decode object identity or surface geometry information that persists different textures. The classifier was evaluated using a leave-one-texture-out cross-validation scheme, in which models were trained on responses to all but one texture and tested on the held-out texture, ensuring that training and testing stimuli were never identical. We performed both 20-way multiclass classification (all objects simultaneously) and pairwise binary classification (all object pairs). The same training and testing time windows were used as in the M2S analysis. This classifier-based approach yielded decoding accuracies highly consistent with the distance-based M2S decoder, confirming that the reported object discriminability is robust to the choice of readout method (Supplemental Figure S3).

### Applying linear decoders to neural responses across time

The choice and lure representations were constructed from a training time window by averaging firing rates across stimulus repetitions, while the sample response was drawn from individual repetitions (trials) within a separate testing time window. Averaging the choice and lure responses yields stable templates and confines single-trial variability to the *sample*, the response of interest. This asymmetry mirrors the behavioral task. The two choice stimuli looped and remained available until a choice was made, whereas the sample was viewed only briefly, so minimizing choice-related uncertainty isolates the reliability of the single-trial sample readout.

We evaluated three training windows (0-100*ms*, 100-300*ms*, and 0-400*ms* relative to stimulus onset) while sliding the 100*ms* testing window in 20*ms* steps from -100*ms* to 600*ms*. All firing rates were latency-adjusted to account for response onset differences across brain regions. Decoding accuracy was computed for each trial by averaging over valid stimulus repetitions, and overall accuracy was obtained by averaging across all object, texture, viewpoint, and size combinations.

### Model zoo

Our model set spans a progression from those derived from image-based supervised learning to those derived from video-based self-supervised learning. CNNs (276 models, 512 total layers) process static images through hierarchical spatial filters, trained with supervised classification objectives prioritizing object categorization. Vision Transformers (ViT, 130 models, 476 total layers) replace convolutions with global attention (queries) over image patches but similarly operate on static frames with supervised training [30]. DINOv2 (4 models, 56 total layers) retains the ViT architecture but removes reliance on labels through self-supervised distillation, where a student network learns to match a teacher network’s representation across augmented views to promote generalization [31]. PerceptionEncoder (6 models, 77 total layers) extends this approach as a large-scale contrastive vision–language model and video–text fine tuning to learn general-purpose embeddings for image and video understanding, though it does not incorporate temporal learning constraints [32].

The remaining model types that we tested depart from the above models as they operate on video and they were trained explicitly on temporal structure. TimeSformer (3 models, 39 total layers) extends transformers to video through factorized spatiotemporal attention, but retains supervised action classification objectives [33]. VideoMAEs (video masked auto-encoders, 10 models, 130 total layers) depart from supervised training and learn spatiotemporal structure by reconstructing masked video patches, forcing the model to internalize regularities of visual features evolving over time [34]. V-JEPAs (4 V-JEPA models, 3 V-JEPA2 models, 223 total layers) advances this approach further by predicting latent representations of masked regions rather than reconstructing pixels, emphasizing abstract spatiotemporal structure over low-level visual details [35, 36]. This progression from supervised image classification through self-supervised image learning, to self-supervised video models that learn predictive regularities in both space and time provides a framework for identifying which computational paradigms lead to models that best approximate the adult neural mechanisms of extracting surface geometry from motion. We refer to V-JEPA and VideoMAE collectively as *predictive coding video models*, as both are trained with self-supervised objectives that require predicting masked spatiotemporal content from visible context in natural videos. We adopt this terminology to emphasize the shared training strategy – learning representations through prediction of withheld sensory content, including future time steps from past context – which we take to be a computationally consequential property linking these models to cortical function; we do not claim architectural correspondence. Our usage is broader than the classical predictive coding framework in neuroscience [37, 38], which posits a hierarchical architecture in which top-down predictions are compared against bottom-up input and prediction errors propagate downstream.

### Extracting features from stimulus-computable models

For each model, we extracted features from the dynamic frames of each stimulus video (12 frames spanning the 100–300 ms motion period of a 60 fps video; onset and offset transients in the first and last 6 static frames were discarded). For image-computable models, features were extracted independently for each frame. For video-computable models, all 12 dynamic frames were provided as a single input and spatiotemporal features were extracted accordingly. When a video model’s expected context length exceeded 12 frames, we symmetrically padded the first and last frames to match the required length. All input preprocessing (pixel normalization, spatial resizing) followed the inference procedures specified by each model’s authors.

Across all architectures, we restricted our analyses to activations from feedforward encoding layers spanning each model’s hierarchy, excluding auxiliary components such as predictor networks in V-JEPA or decoder networks in VideoMAE. To obtain a common representational format, we applied a two-stage dimensionality reduction pipeline uniformly across models. In the first stage, PCA (250 components) was fit on features extracted from an independent set of shape-from-motion stimuli comprising different object geometries and surface textures than those used in the main experiment. In the second stage, the learned PCA projections were applied to features from all experimental videos. This projection preserved a substantial amount of geometric structure of frame-by-frame time series (see Supplement S6 for further details). Features were extracted across a range of layers spanning each model’s full hierarchy to enable layer-resolved comparisons with neural data.

### Encoding analyses

For univariate prediction, we fit a PLS model (25 components) per neuron on five of six texture classes and 18 of 20 objects, evaluating on the held-out texture and objects. Predictivity was the correlation between predicted and observed spike counts integrated over the dynamic period (100–300 ms), normalized by each neuron’s split-half reliability (noise ceiling) and accounting for region-specific response latencies. CKA mapping followed the procedure in Supplement S5.2. Additional details for both univariate and multivariate mapping and prediction are described in Supplement S5.

## Acknowledgements

We are grateful to Alexis Mackiewicz, Amanda Armijo, Natalie Gutierrez, and Emily Zhang for their expert veterinary and surgical collaboration; to Sarah Goulding and Alina Peter for their dedicated animal care and surgical support; and to Carolyn Wan-hsun Wu for her technical guidance with histology. Neural recording hardware was supported by Modular Bionics (N-form arrays), Blackrock Neurotech (Utah arrays), CortexPlore (precision surgical guidance), and ActuatedNeuroscience (NeuralGlider). Computational resources were provided by MIT’s Office of Research Computing and Data (ORCD). This work was supported by Office of Naval Research (ONR) Award N000142112801.

## Declarations

### Funding

This work was supported by the Office of Naval Research (MURI Award N000142112801 to James DiCarlo and Joshua Tenenbaum), the National Science Foundation (Award 2124136), and the Simons Foundation (Awards 542965 and NC-GB-CULM-00002986-04).

### Conflict of interest/Competing interests

James DiCarlo serves on the Yale University’s Wu Tsai Institute Advisory Board, the External Advisory Board of the AI Institute for Artificial and Natural Intelligence (ARNI), and the Advisory Committee of the Lefler Center at Harvard Medical School. The remaining authors declare no competing interests.

### Ethics approval and consent to participate

All procedures involving non-human primates were performed in accordance with the National Institutes of Health Guide for the Care and Use of Laboratory Animals and were approved by the Massachusetts Institute of Technology Committee on Animal Care (CAC protocol no. 2509000846A001). Animals were housed and maintained by the MIT Department of Comparative Medicine (DCM). All behavioral experiments involving human participants were conducted in accordance with the MIT Committee on the Use of Humans as Experimental Subjects (COUHES), protocol no. 0812003043. Informed consent was obtained from all human participants prior to participation.

### Consent for publication

Not applicable.

### Data availability

The neural data supporting the findings of this study will be made publicly available upon publication at DANDI archive (dandiarchive.org).

### Materials availability

Stimulus videos and experimental materials will be made available upon publication at DANDI archive (dandiarchive.org). Custom neural recording hardware specifications are described in Methods and available upon request.

### Code availability

Source code will be made available upon reasonable request to the corresponding author.

### Author contribution

YB, TPO, JBT, and JD conceived of and designed the experiments. TPO and YF created the stimuli and conducted the behavioral experiments. YB conducted the neuroscience experiments, developed a novel surgical protocol for chronic neural array implantation in deep parietal cortex, and performed neural recordings. AAH assisted with neural recordings and surgeries. HM conducted histology analyses. TPO and YB implemented the computational models and analyses. YB, TPO, JBT, and JD interpreted the results. YB wrote the paper with input from TPO, YF, JBT, and JD.

## Supplementary information

### S1 Recording sites within the inferior parietal lobule (IPL)

We targeted the inferior parietal lobule (IPL) as a downstream stage of the dorsal visual pathway, selected to parallel the role of IT cortex in the ventral pathway. As with IT cortex, where we recorded from multiple subregions (aIT, cIT, pIT), we sampled two areas within the IPL: area Tpt (temporo-parietal) and area 7op (parietal operculum). Area 7op is situated near the fundus where the intraparietal sulcus (IPS) and lateral sulcus converge [24, 25], and occupies a relatively more downstream position within the IPL hierarchy compared to area Tpt [49]. Recording site locations were verified through structural MRI acquired in vivo and confirmed post-mortem via histological analysis (Fig. S1).

Following completion of research in one of the animals involved in dorsal stream recording, the decision to proceed with perfusion and histological analysis was made in consultation with the attending veterinarian from the MIT Department of Comparative Medicine (DCM). Together, we determined that euthanasia was the most humane course of action given the animal’s advanced age and naturally declining condition. All procedures were conducted in accordance with protocols approved by the MIT Committee on Animal Care (CAC). The animal was deeply anesthetized under veterinary supervision and euthanized with an intravenous dosage of sodium pentobarbital (100–120 mg/kg). Euthanasia was confirmed by cessation of heartbeat and respiration via auscultation and absence of withdrawal reflex to firm toe pinch. The animal was then perfused transcardially, first with phosphate-buffered saline (PBS) to clear the vasculature, followed by a cold solution of 4% paraformaldehyde (PFA) in PBS. The brain was extracted and post-fixed in 4% PFA for approximately ten days to ensure complete fixative penetration, then cryoprotected by sequential immersion in 4% PFA with increasing sucrose concentrations (10% to 30%). Upon removing the neural recording device, we sectioned the right hemisphere horizontally on a freezing microtome at 60 micrometer thickness. Sections were collected into sodium azide solution for storage. Every third and sixth section was stained for Nissl substance using cresyl violet to delineate cytoarchitectonic boundaries of cortical areas and laminar organization. Stained sections were cover-slipped and imaged on an epifluorescence microscope (Zeiss Axio Imager Z2) to identify electrode tract locations from the N-Form array shanks across cortical layers.

**Fig. S1.**
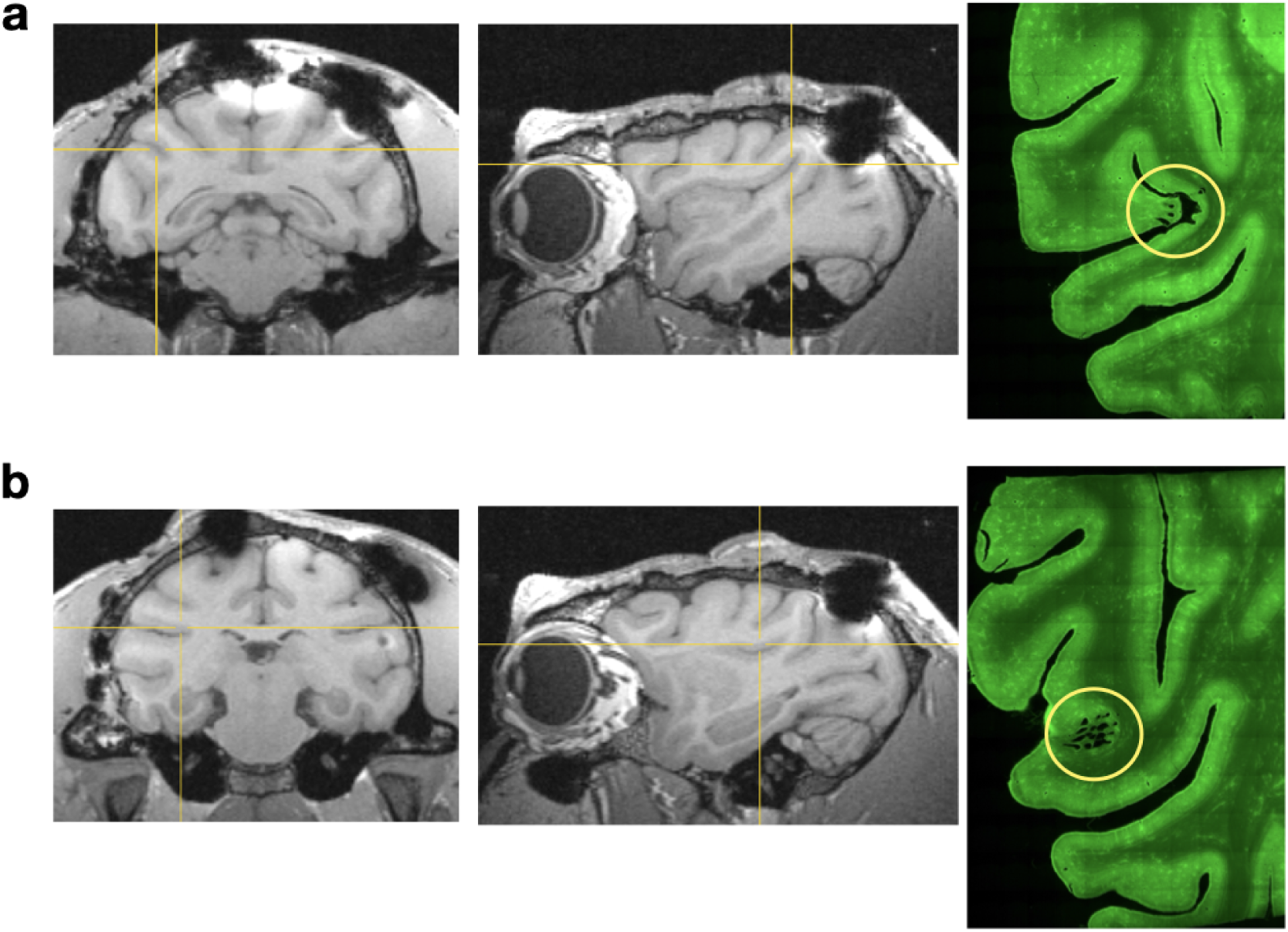
Structural anatomy of target recording sites in the dorsal visual pathway. **(a) Area Tpt (temporo-parietal area).** Left and center panels show structural MRI sections (coronal and sagittal, respectively) from the first monkey used for dorsal recordings, centered on area Tpt (thin crosshairs). Large, dark shadows on the skull correspond to cranial implants which distorted local imaging regions. **Right panel** shows a matched histological section obtained post-mortem from the same animal. The yellow circle indicates the sulcal bank where the intraparietal sulcus and lateral sulcus converge, corresponding to the region where array shanks were positioned. Small dark perforations mark individual shank penetration sites. **(b) Area 7op (parietal operculum).** A subset of array shanks were elongated to reach the fundus of the parietal operculum. Same conventions as in (a).

### S2 Receptive fields estimated from each target region

Receptive fields (RFs) were mapped for each recorded unit using a moving bar paradigm adapted from Viswanathan & Nieder (2017) [50]. During fixation, a bar stimulus was presented sequentially at each location in a grid spanning 42°x24° of visual space (15 horizontal positions from -21° to 21° and 9 vertical positions from -12° to 12°, at 3° intervals), covering 135 unique locations. At each grid position, the bar was swept across the location along multiple directions and orientations; the evoked spiking response at each position was computed by integrating spike counts over the stimulus presentation window after stimulus onset, with respect to the neural unit’s response latency. The mean firing rate across repetitions at each grid location produced a two-dimensional response map for each unit. To estimate RF boundaries, a two-dimensional Gaussian function was fitted to each unit’s response map, and the fit quality was assessed by the Pearson correlation between the fitted and observed maps. Units whose Gaussian fits exceeded a minimum correlation threshold (0.5) were plotted in Figure S2. For visualization, the fitted Gaussian was then upsampled to native stimulus resolution (approximately 50 pixels/°) and thresholded at 1 standard deviation (corresponding to 60% of peak amplitude) to generate a binary RF mask for each unit. Individual unit masks were cropped to the stimulus field of view and superimposed to produce a population-level RF coverage map for each recording region. RF centers are concentrated in the contralateral visual field, consistent with known retinotopic biases in both pathways.

**Fig. S2.**
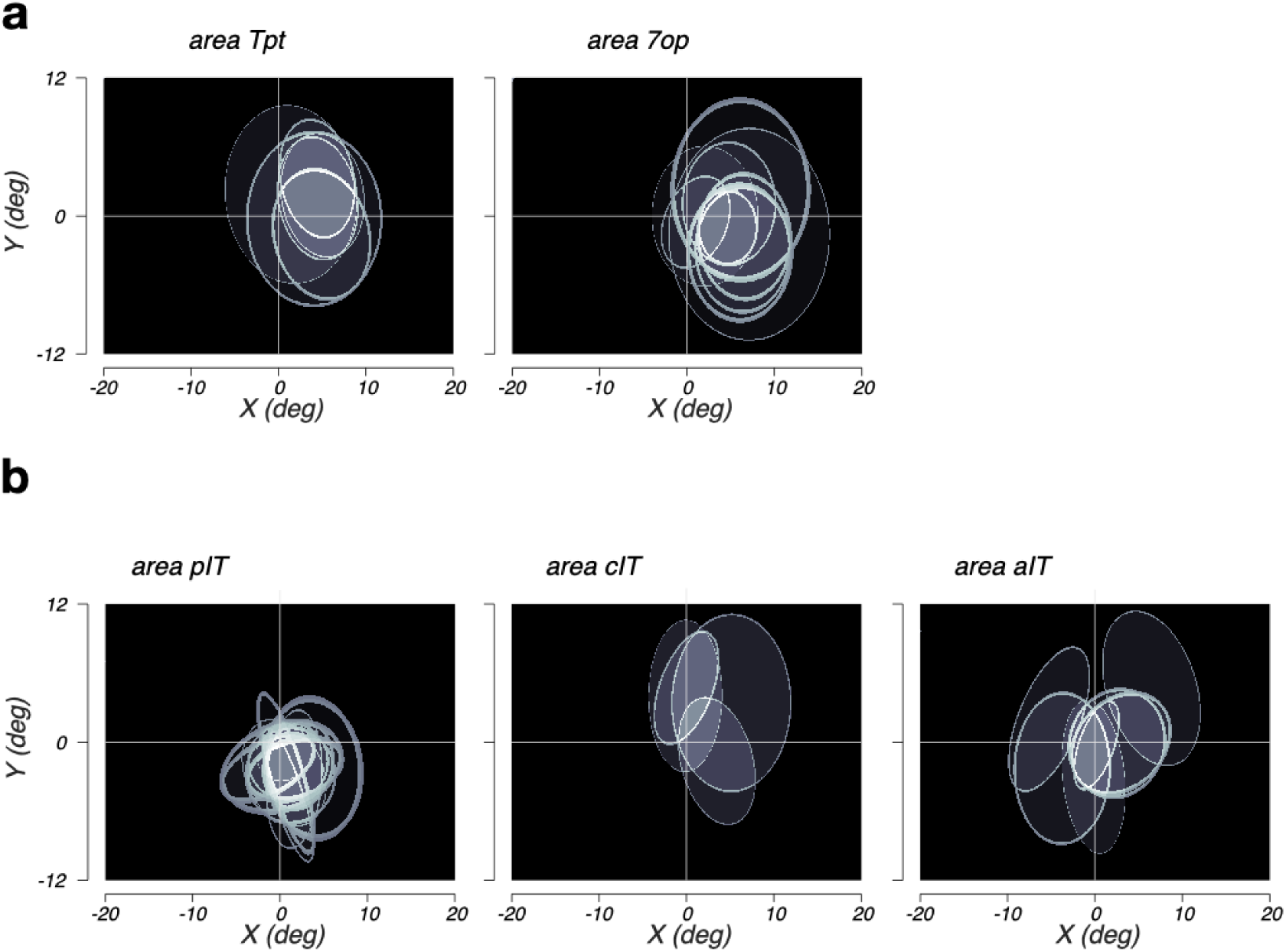
Receptive fields from downstream regions in the dorsal and ventral visual pathways. **(a) Inferior Parietal Lobule (downstream in dorsal stream)**. RF coverage for a subset of IPL units recorded from left hemisphere in one animal (N = 12, and 8 units across areas 7op and Tpt). **(b) Inferior Temporal Cortex (downstream in ventral stream)**. RF coverage for a subset of IT cortex units from the left hemisphere in one animal (N = 7, 4, and 15 units across aIT, cIT, and pIT).

### S3 Decoding spiking activity using logistic regression classification

As an alternative to the distance-based match-to-sample decoder, we trained a single L2-penalized multinomial logistic regression to classify the 20 objects in the stimulus set.

Generalization across nuisance variables, such as texture class, was enforced by a leave-one-texture-out cross-validation policy. For each fold, the classifier was fit on trials from all but one surface texture and evaluated on the held-out texture, cycling through all textures. In Figure 3 and in Supplemental Figure S3, spikes rates estimated within the 100-300ms window (latency-adjusted) served as the training signal. Using this time window, training data were pooled across all three stimulus sizes (6^0^, 16^0^, and 27^0^). A single decoder was fit per texture-fold on the size-pooled set and then evaluated separately at each size, so that the size-invariance of the readout could be measured rather than designed in. Features were z-scored across the training set before fitting. Training and test trials were additionally restricted to the same viewpoint set, so that the decoder had to generalize only across textures and trial repeats. To plot read-outs across time, test-trial responses were read out in 100-ms windows stepped by 20 ms from -100 to +600 ms relative to stimulus onset.

Although the classifier itself is a 20-way multiclass model, we converted its outputs to binary 2AFC reports to match the behavioral task. For each held-out test trial, we extracted the full class-posterior vector, selected the probabilities assigned to the designated sample and lure objects, and re-normalized them to sum to one: *p*^(sample) = p(sample) / (p(sample) + p(lure)). The trial was scored correct if *p*^(sample) *>* 0.5, i.e., if the decoder assigned greater probability to the true object than to the lure. This procedure yields, for every ordered [sample, lure] object pair, a pairwise accuracy derived from a single shared multiclass fit rather than from separately trained binary classifiers.

**Fig. S3.**
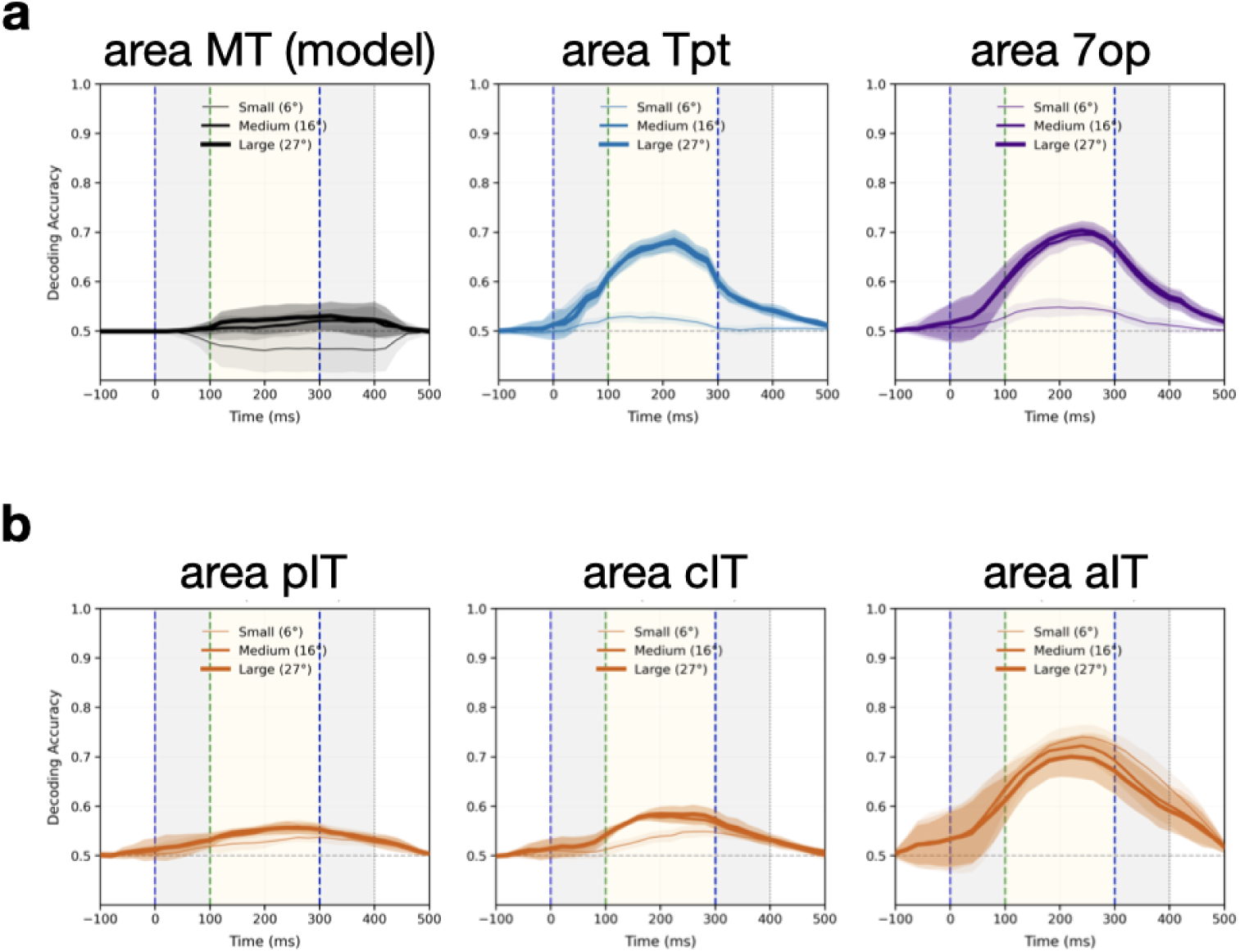
Match-to-sample read outs from dorsal and ventral visual pathways. **(a) Regions from the dorsal visual pathway (downstream left-to-right). (b) Regions from the ventral visual pathway (downstream left-to-right).**

### S4 Partial Least Squares (PLS) regression-based mapping

PLS regression discovers a low-dimensional basis by iteratively extracting components that maximize covariance between feature and response spaces. The resulting weight matrix *W_P LS_* = [*w*_1_*, w*_2_*, …, w_k_*] consists of basis vectors ordered by their contribution to explained variance, where each component *w_i_*is orthogonal by design. This supervised dimensionality reduction explicitly targets variance relevant to the observed neurons, typically achieving high prediction accuracy with relatively few components. The learned basis effectively filters out feature dimensions uncorrelated with neural responses, implementing an efficient compression that prioritizes neuron-by-neuron predictive power.

### S5 Univariate and multivariate neural mapping

**Fig. S4.**
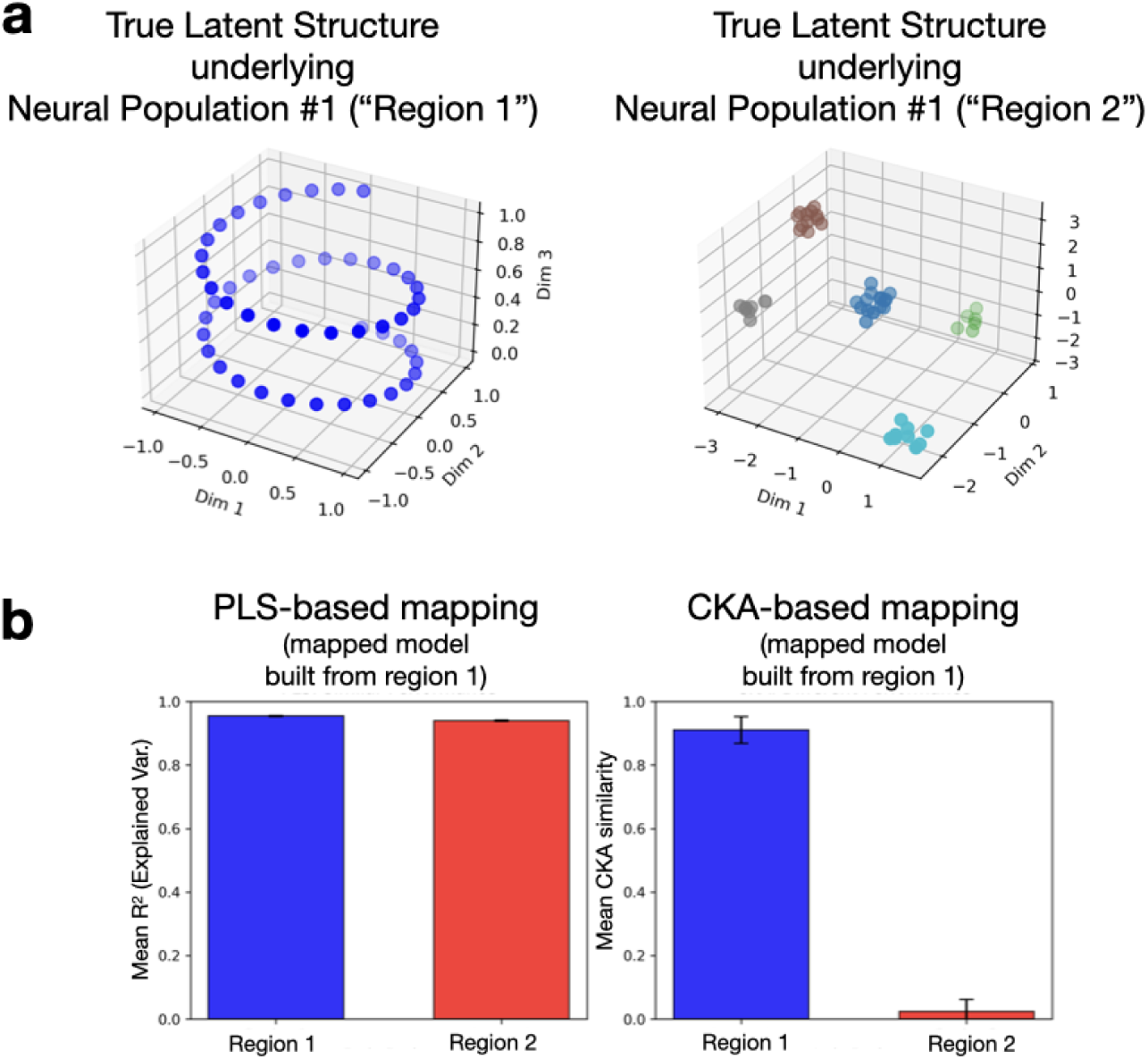
A simulation in which two neural populations can have similar explained variance while having fundamentally different coordinated activity, or geometric characteristics, in their representation space. (a) Two neural populations were simulated from distinct latent structures. The left figure illustrates a latent low-dimensional trajectory used for generating spike rates for Population 1 (region 1). The right-hand side figure shows a discrete clustered structure used for generating spike rates for Population 2. For each region, the latent structure was projected to 100 units and the mean spike rate was matched between the two regions. (b) Surrogate model features (e.g., from a vision model) were designed to reflect population 1 (region 1), but not population 2. (Left figure) Model features were mapped to individual units in regions 1 and 2 using PLS-regression (25 components). (Right figure) Model features were mapped by optimizing separate weight matrices for each region 1 and 2. Geometry based mapping distinguishes the two regions whereas PLS-based mapping (maximizing explained-variance per-unit) does not. Geometry-based signals are better distinguished with CKA-based mapping as the true latent structure in each region is different.

#### S5.1 Partial Least Squares (PLS) regression-based mapping

PLS regression discovers a low-dimensional basis by iteratively extracting components that maximize covariance between feature and response spaces. The resulting weight matrix *W_P LS_* = [*w*_1_*, w*_2_*, …, w_k_*] consists of basis vectors ordered by their contribution to explained variance, where each component *w_i_*is orthogonal by design. This supervised dimensionality reduction explicitly targets variance relevant to the observed neurons, typically achieving high prediction accuracy with relatively few components. The learned basis effectively filters out feature dimensions uncorrelated with neural responses, implementing an efficient compression that prioritizes neuron-by-neuron predictive power.

#### S5.2 Centered Kernal Alignment (CKA)-optimized mapping

PLS-based mapping (unit-wise explained-variance) alone is insufficient for characterizing geometric properties embedded in coordinated activity. For example, populations can achieve similar explained variance across neurons (average explained-variance), while exhibiting fundamentally different population-level geometric properties ([41]; also see Supplemental Figure S4). This is because PLS-regression finds linear combinations of model response to maximize variance explained across stimulus conditions, but does not necessarily preserve geometric signal embedded in population activity as the learned basis for mapping units are not rotation-invariant – in this scenario, geometrical similarities from linear regression is an artifact, not an objective.

On the other hand, a CKA-based optimization approach instead finds a linear mapping *W* by maximizing the alignment between mapped artificial neural network features (*XW*) and biological neural responses (*Y*), where the optimization criterion is based on geometrical similarity, such as *CKA*(*XW, Y*). Using debiased linear CKA [51] to handle the high-dimensional, low-sample regime typical of neuroscience data, we optimize for a weight matrix *W_CKA_* that preserves the inner product structure between all pairs of neural population responses. This approach maintains pairwise distances and angles between activity patterns across conditions or stimuli. Unlike PLS, this method is unsupervised with respect to individual neuron identities—it preserves the geometric structure of population responses rather than targeting specific neural predictions. The resulting basis typically produces denser mappings than PLS solutions, as it must maintain all geometric relationships rather than just those relevant for prediction. The distinct algebraic structures above reflect a fundamental trade-off in neural system identification, particularly relevant when analyzing different response characteristics, e.g., visual versus motor cortices [3, 41, 52]. While regression-based mapping produces sparse, low-rank mappings optimal for predicting individual neural responses, geometry-based mapping preserves the relational structure determining how neural populations distinguish between stimulus—wise pairwise distances, angles, and manifold geometry that may be critical for neural readouts in downstream areas of a brain region. Empirical comparisons using split-half reliability as noise ceilings reveal that while PLS may achieve higher explained variance for individual neurons, CKA-based mappings better preserve coordinated activity within a neural population (correlation structure). The choice between these approaches should align with the scientific question: PLS for maximizing prediction accuracy of individual unit responses, CKA-based optimization for discovering transformations that preserve the geometric structure underlying neural computation.

The dissociation between unit-level prediction accuracy and population-level geometric structure parallels recent findings in the neural network interpretability literature. Melander and colleagues demonstrate that hidden units with similar activation magnitudes can exert fundamentally different causal contributions to network output, and that decomposing these contributions at the population level reveals computational motifs invisible to unit-wise analysis [53]. Together, these observations reinforce that explained variance for individual neurons is necessary but insufficient for characterizing the representational correspondence between artificial and biological systems — motivating the geometry-preserving mapping approach adopted here.

**Fig. S5.**
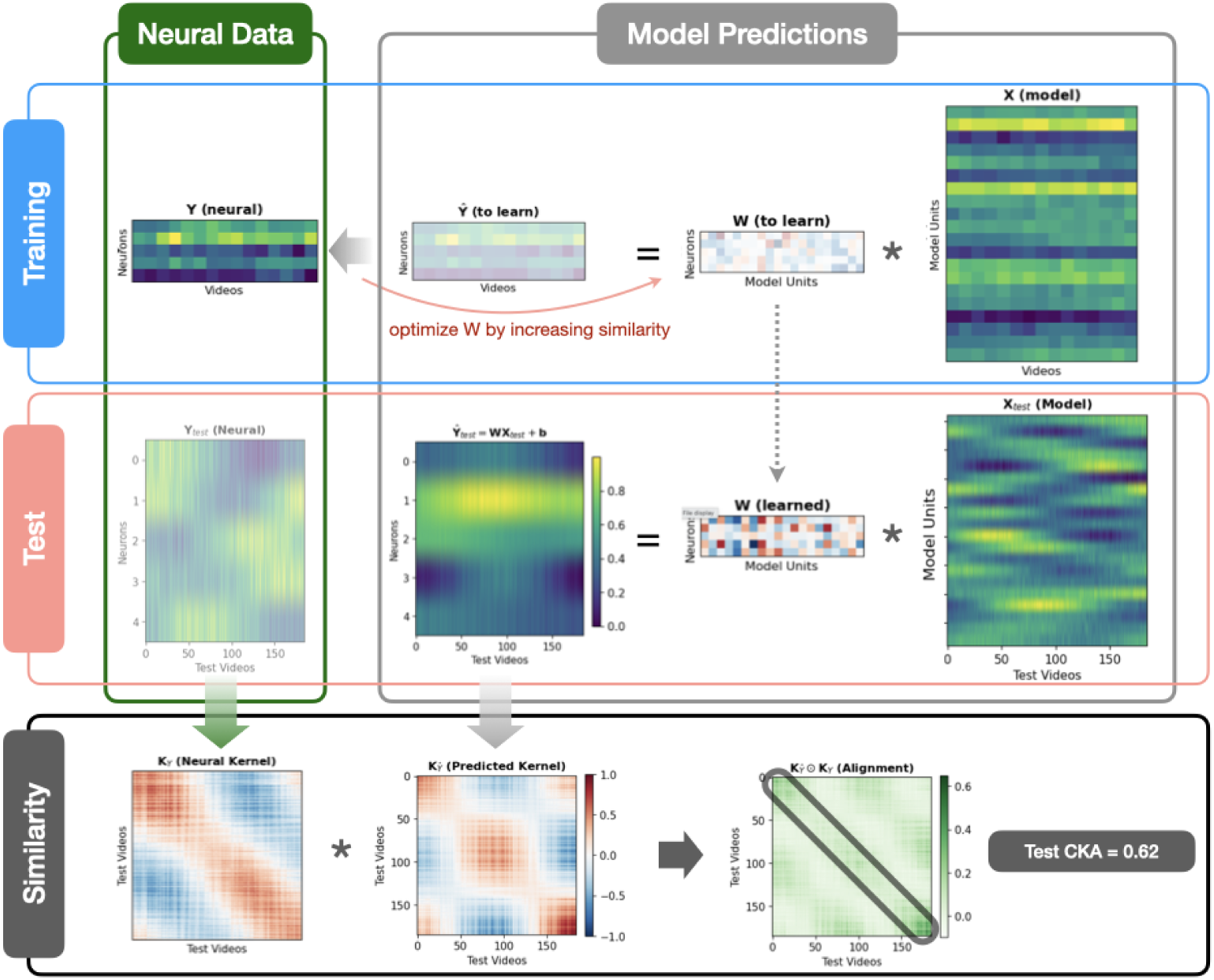
Pipeline for cross validating CKA similarities between neural data (spiking activity) and model activations.

**Fig. S6.**
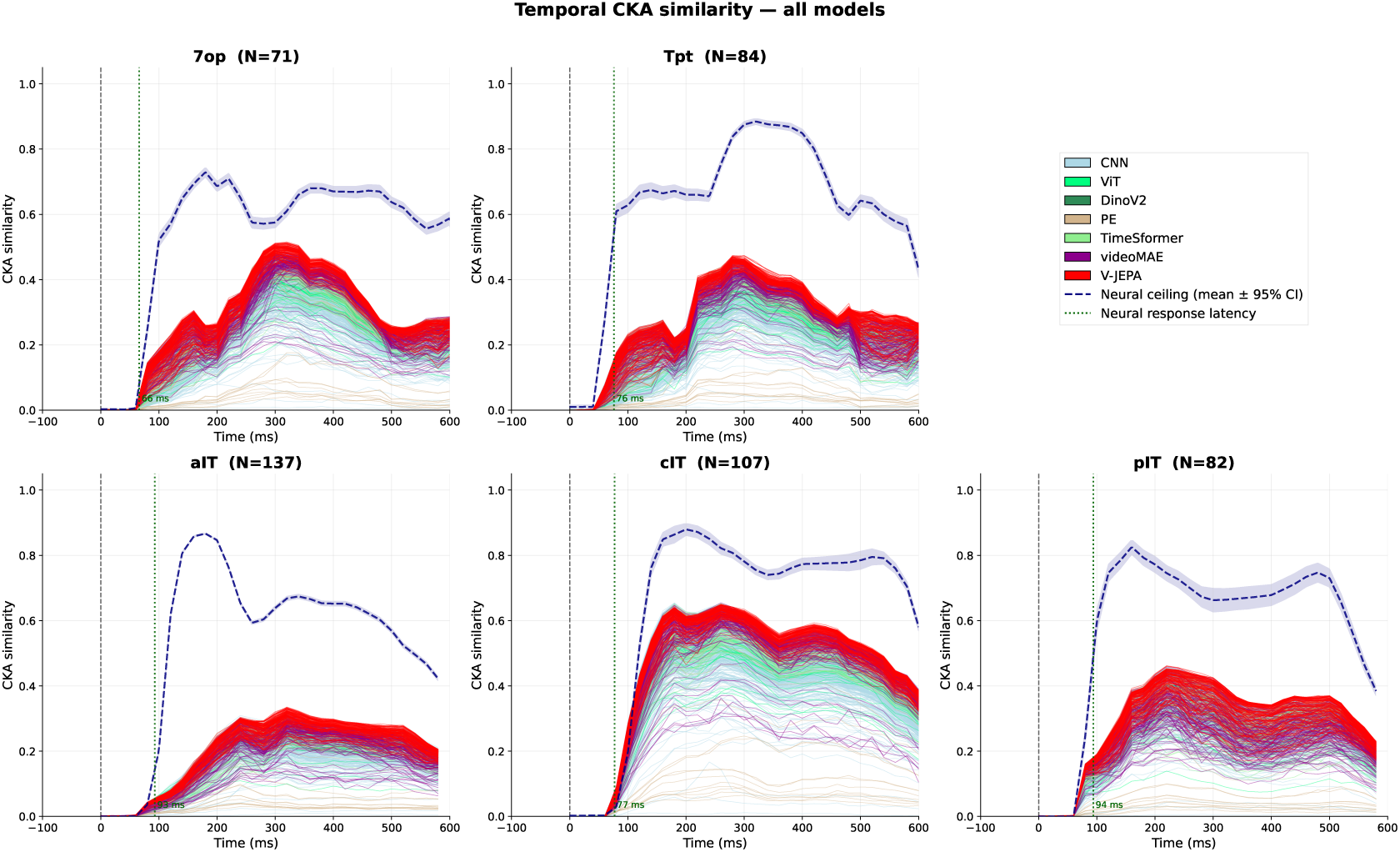
Temporal CKA similarity between neural population responses and computational model representations. Each panel shows one cortical region; IPL (downstream dorsal) regions in top row and IT cortex in bottom row (N = number of units per region). Thin lines show the debiased linear CKA similarity as a function of time for individual model layers, colored by model family (CNN, ViT, DinoV2, PerceptionEncoder, TimeSformer, videoMAE, V-JEPA). The dashed navy line with shaded band shows the mean neural reliability ceiling ± 95% CI across all models, estimated via split-half CKA (described below). The dotted green vertical line marks the region’s mean neural response latency (rounded up to the nearest ms). The dashed black vertical line at 0 ms marks stimulus onset. **CKA timeseries computation.** For each region, spike count responses were integrated over a 100 ms sliding window stepped in 20 ms increments, spanning -100 to 600 ms relative to stimulus onset. Neural responses were shifted by the region’s population response latency prior to analysis, so that time bins are expressed in the model’s reference frame, i.e., time 0 corresponds to stimulus onset for the model. At each timepoint, A linear mapping from model features to neural population responses was trained to maximize CKA similarity (debiased linear CKA) on held-out stimuli via 6-fold cross-validation stratified by texture class. The neural reliability ceiling was estimated by computing split-half CKA between two random halves of neural repeats.

### S6 PCA-based dimensionality reduction of model features

**Fig. S7.**
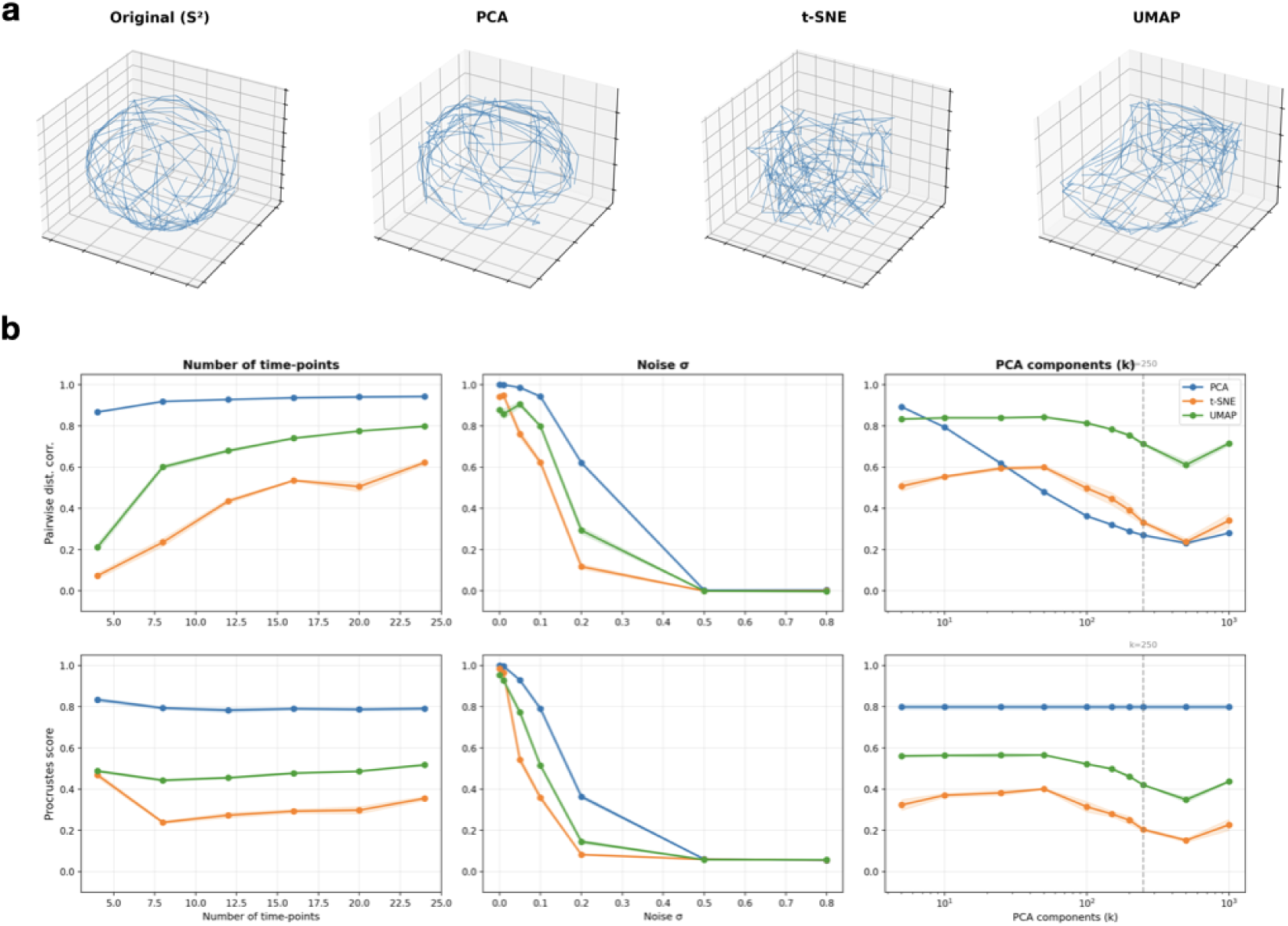
**(a) Simulation of dimensionality-reduced time series data.** Geodesic trajectories simulated on a 2-sphere (*S*^2^) shown in the original high-dimensional space (left) and after projection by PCA, t-SNE, and UMAP (right three panels). Each panel displays the same set of random-walk trajectories; visual comparison reveals how each method distorts or preserves the global topology of the spherical manifold. **(b) Summary of geometrical distortions from simulated time series data** Geometric structure preservation as a function of three parameters (columns): number of time points, noise level (*σ*), and number of PCA components (k). Two metrics are shown (rows): pairwise distance correlation (top) and Procrustes score (bottom). Lines compare PCA (blue), t-SNE (orange), and UMAP (green). Higher values indicate better preservation of the original manifold geometry. In the rightmost column, the vertical dashed line indicates the reduced dimensionality (*k* = 250) applied to high-dimensional model features.

To compare model feature dynamics with neural population responses across time, we first reduce the dimensionality of model features using principal component analysis (PCA). The choice of retaining the first 250 components may appear arbitrary; here we provide a simulation-based justification (Supplemental Figure S7).

Model features extracted from video foundation models (e.g., V-JEPA2) can occupy feature spaces with millions of dimensions, far exceeding the number of simultaneously recorded neurons. To bring these representations into a commensurate dimensionality while preserving as much geometric structure as possible, we considered widely-used dimensionality reduction methods to seek which method might lead to a low-dimensional trajectory that captures dominant structure of coordinated activity across many features.

To assess how well this procedure preserves temporal geometric structure, we conducted a series of controlled simulations. We generated random geodesic trajectories on the 2-sphere (*S*^2^ for mathematical convenience), providing ground-truth structure with known geometry. Each 3-D time point was embedded into a high dimensional space (e.g., 100,000) via a random orthogonal projection with additive Gaussian noise (e.g., *σ* = 0.1), mimicking the high-dimensional aspect of model features. We then applied PCA, t-SNE, and UMAP to features at each time point, reducing back to a low-dimensional space while keeping the same number of time points, exactly as in our analysis pipeline. We quantified geometric fidelity using two metrics: (1) Spearman correlation between pairwise distances in the original and reduced spaces, and (2) Procrustes alignment score between the original and reduced trajectories (after optimal rotation, scaling, and translation).

We swept across the number of retained PCA components (k = 5, 10, … 1,000). Both metrics improved rapidly with increasing k and began to plateau around k = 100–250. At k = 250, PCA achieved high pairwise distance correlation and Procrustes scores, substantially outperforming t-SNE and UMAP. Geometric fidelity was robust to the number of time points examined and degraded only at high noise levels (*σ >* 0.3), consistent with a signal-to-noise regime well within what is expected for pretrained model features (even with moderate dropout and batch-normalization stochasticity; we disabled dropout during model evaluation).

Taken together, these simulations indicate that retaining 250 PCA components strikes a practical balance. It preserves the dominant geometry of high-dimensional feature trajectories, reduces dimensionality by several orders of magnitude, and yields a feature space that is commensurate in size with recorded neural populations. We therefore use k = 250 throughout our analyses.

### S7 Predictions from synthetic populations derived by each model encoding layer

**Fig. S8.**
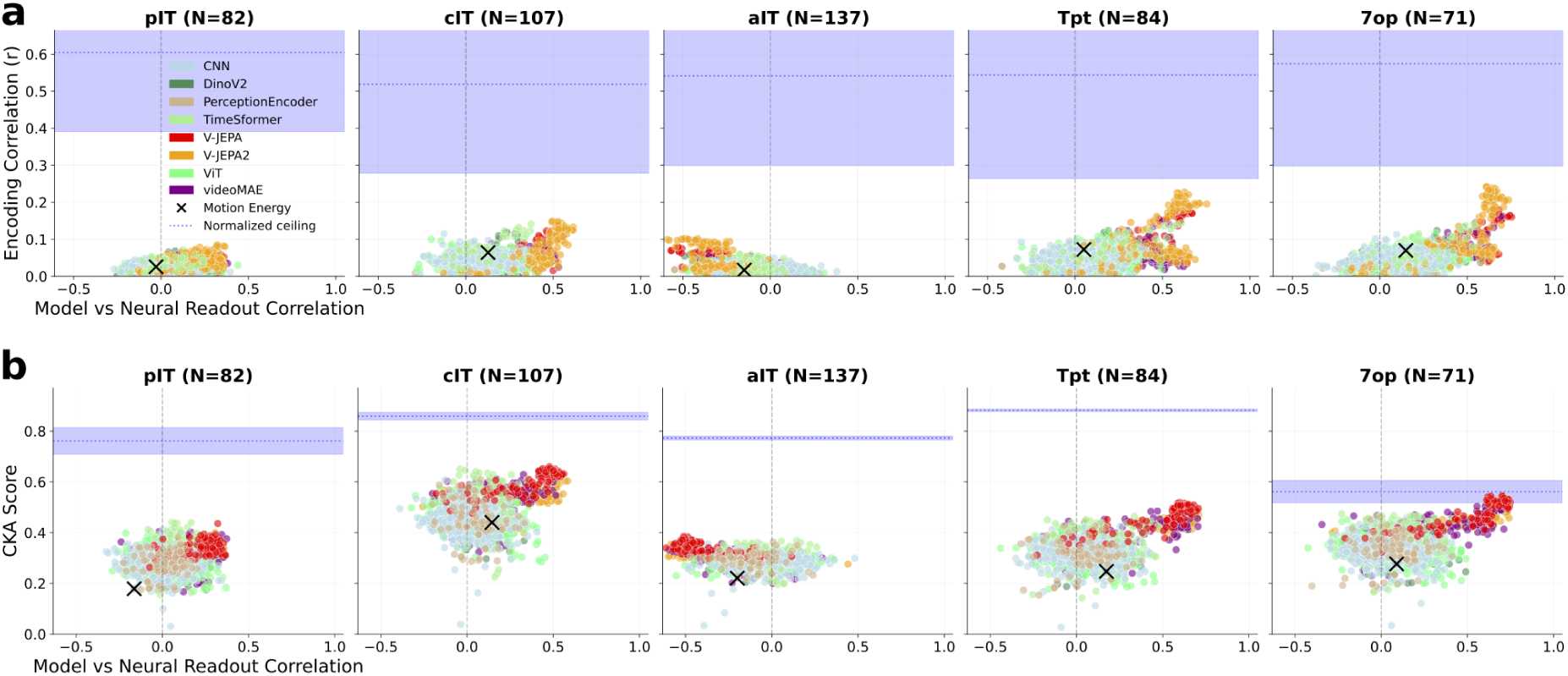
Generalizing neural encoding and behavioral read-outs for synthetic neural populations across all model layer activations. Raw data points indicate surrogate populations derived from each model layer in each model class, which was summarized in Figure 6, (a) Synthetic population generated unit-by-unit from stimulus-computable models (using PLS regression). Each panel corresponds to a cortical region (same with Figure 6. Each point represents the best model layer from each model, color-coded by its model class. The x-axis shows the readout similarity between model neurons and biological neurons–defined by the Spearman correlation coefficient calculated from readout accuracies across the same held-out conditions. The y-axis shows the encoding correlation between PLS-mapped synthetic population responses and recorded neural responses, averaged across held-out conditions. The blue horizontal dashed line and shaded region indicate the noise ceiling of PLS predictivity in the recorded data (mean split-half reliability *±* SD across units). (b) Synthetic neurons generated at the level of populations by matching coordinated activity across neurons (CKA similarity). Same plotting conventions as (a). The blue horizontal dashed line and shaded region indicate the noise ceiling of CKA scores (mean and standard deviation from split-half resamples).

## Footnotes

1 PerceptionEncoder was trained on 5.4 billion image–text pairs (MetaCLIP; 86 billion total samples seen) with 22 million additional video–text finetuning samples, exceeding the data scale of all other models evaluated.

## Notes

### Summary of Updates

Introduction updated to clarify problem statement.

